# Functional monovalency amplifies the pathogenicity of anti-MuSK IgG4 in myasthenia gravis

**DOI:** 10.1101/2020.09.24.296293

**Authors:** Dana L.E. Vergoossen, Jaap J. Plomp, Christoph Gstöttner, Yvonne E. Fillié-Grijpma, Roy Augustinus, Robyn Verpalen, Manfred Wuhrer, Paul W.H.I. Parren, Elena Dominguez-Vega, Silvère M. van der Maarel, Jan J. Verschuuren, Maartje G. Huijbers

## Abstract

Human IgG4 usually displays anti-inflammatory activity, and observations of IgG4 autoantibodies causing severe autoimmune disorders are therefore poorly understood. In blood, IgG4 antibodies naturally engage in a stochastic process termed Fab-arm exchange in which unrelated IgG4s exchange half-molecules continuously. The resulting IgG4 antibodies are composed of two different binding sites, thereby acquiring monovalent binding and inability to cross-link for each antigen recognized. Here, we demonstrate this process amplifies autoantibody pathogenicity in a classic IgG4-mediated autoimmune disease: muscle-specific kinase (MuSK) myasthenia gravis (MG). In mice, monovalent anti-MuSK IgG4s caused rapid and severe myasthenic muscle weakness, whereas the same antibodies in their parental bivalent form were less potent or did not induce a phenotype. Mechanistically this could be explained by opposing effects on MuSK signaling. Isotype switching to IgG4 in an autoimmune response thereby may be a critical step in the development of disease. Our study establishes functional monovalency as a novel pathogenic mechanism in IgG4-mediated autoimmune disease and potentially other disorders.

**Graphical abstract:** 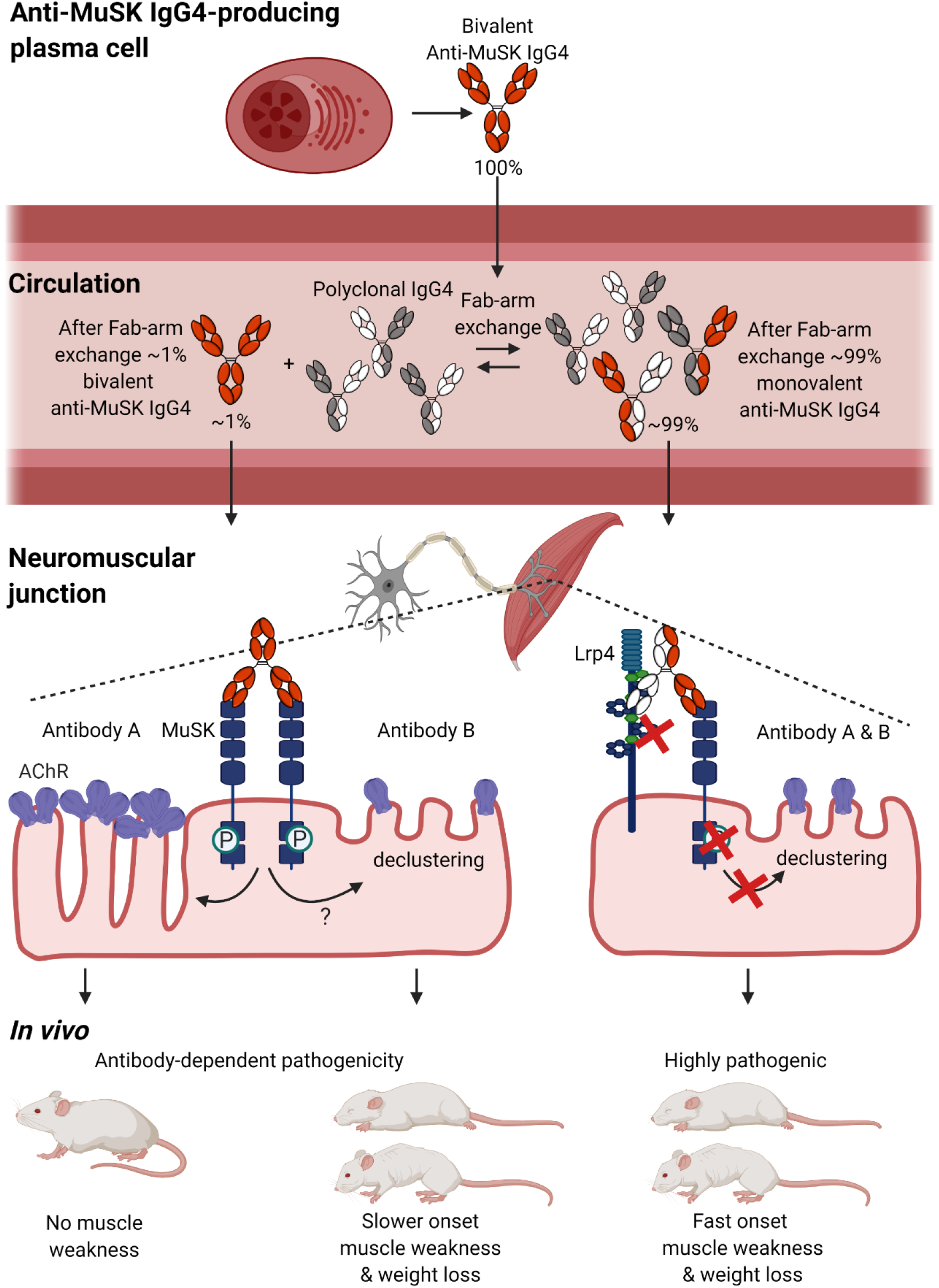

## Introduction

Recently, a new and growing class of antibody-mediated autoimmune diseases characterized by predominant pathogenic IgG4 responses was described (Huijbers et al., 2018; Huijbers et al., 2015; Koneczny, 2018). IgG4 is a peculiar antibody with unique characteristics. It is for example unable to activate complement and has low affinity for Fcγ receptors on immune cells (Lighaam and Rispens, 2016; Vidarsson et al., 2014). It is therefore considered anti-inflammatory and the pathogenicity of IgG4 autoantibodies is sometimes questioned. IgG4 molecules furthermore have the unique ability to stochastically exchange half-molecules with other IgG4s in a dynamic process called Fab-arm exchange (Van Der Neut Kolfschoten et al., 2007). This is an efficient process resulting in the vast majority of IgG4 molecules in circulation being bispecific and functionally monovalent for each antigen recognized. Whether the unique functional characteristics of IgG4 (like Fab-arm exchange) influence their pathogenicity in IgG4 autoimmune diseases is not known.

Myasthenia gravis (MG) with antibodies against muscle-specific kinase (MuSK) is one of the first recognized IgG4-mediated autoimmune diseases. MuSK autoantibodies are predominantly of the IgG4 subclass; although anti-MuSK IgG1 and IgG3 may be present concurrently at lower titers (McConville et al., 2004). Anti-MuSK IgG4s induce MG in a dose-dependent manner both in patients and in mice (Klooster et al., 2012; Niks et al., 2008). MuSK, a receptor tyrosine kinase, has a crucial role in establishing and maintaining neuromuscular junctions (NMJs) by orchestrating postsynaptic acetylcholine receptor (AChR) clustering, which is critical for neurotransmission (Burden et al., 2013). Most MuSK autoantibodies bind the extracellular N-terminal Ig-like 1 domain and thereby block the activation of MuSK by low-density lipoprotein receptor-related protein 4 (Lrp4) and agrin (Hoch et al., 2001; Huijbers et al., 2016; Huijbers et al., 2013; Koneczny et al., 2013; McConville et al., 2004; Otsuka et al., 2015). This eventually leads to disassembly of densely packed AChRs in the NMJ, failure of neurotransmission and consequently muscle weakness (Cole et al., 2008; Ghazanfari et al., 2014; Klooster et al., 2012; Morsch et al., 2012; Viegas et al., 2012). *In vitro* characterization of monoclonal antibodies derived from MuSK MG patients furthermore suggests that the valency of MuSK antibodies determines their effects on MuSK signaling (Fichtner et al., 2020; Huijbers et al., 2019b; Takata et al., 2019). Monovalent Fab fragments recapitulate the inhibitory effects of patient-purified IgG4 on MuSK signaling *in vitro* (Fichtner et al., 2020; Huijbers et al., 2019b; Huijbers et al., 2013; Koneczny et al., 2013). Surprisingly, monospecific bivalent MuSK antibodies acted oppositely as (partial) agonists (Huijbers et al., 2019b). To investigate whether IgG4 predominance is critical for disease development in IgG4-mediated autoimmunity and study the role of Fab-arm exchange and autoantibody valency, we generated stable bispecific functionally monovalent MuSK antibodies and their monospecific bivalent equivalents and assessed their pathogenicity in NOD/SCID mice.

## Results

### Generation of patient-derived stable bispecific monovalent IgG4 MuSK antibodies

To investigate the role of MuSK antibody valency on their pathogenicity *in vivo*, pure and stable bispecific MuSK antibodies are needed. Remaining monospecific MuSK antibodies may have confounding effects, e.g. by competing for binding with MuSK and thereby masking effects of the bispecific MuSK antibodies. We therefore adapted the controlled Fab-arm exchange (cFAE) method to IgG4 (Labrijn et al., 2013; Labrijn et al., 2014). In addition to two previously identified patient-derived recombinant MuSK antibodies (13-3B5 and 11-3F6) (Huijbers et al., 2019b), the b12 antibody was chosen to generate an innocuous arm in the bispecific MuSK antibody, because its antigen (the HIV-1 envelope gp120) does not exist in the model systems used (Barbas et al., 1993). To stabilize the antibodies under physiological conditions, the serine (S) at amino acid position 228 was converted into a proline (P) in the IgG4 heavy chain of the anti-MuSK antibodies (Fig. 1A). The b12 antibody was made suitable for efficient cFAE by altering S228P, F405L and R409K in its IgG4 heavy chain. For clarity, the parental monospecific IgG4 will be designated as bivalent, and the bispecific IgG4 as monovalent towards MuSK in the remainder of the manuscript.

**Fig. 1.**
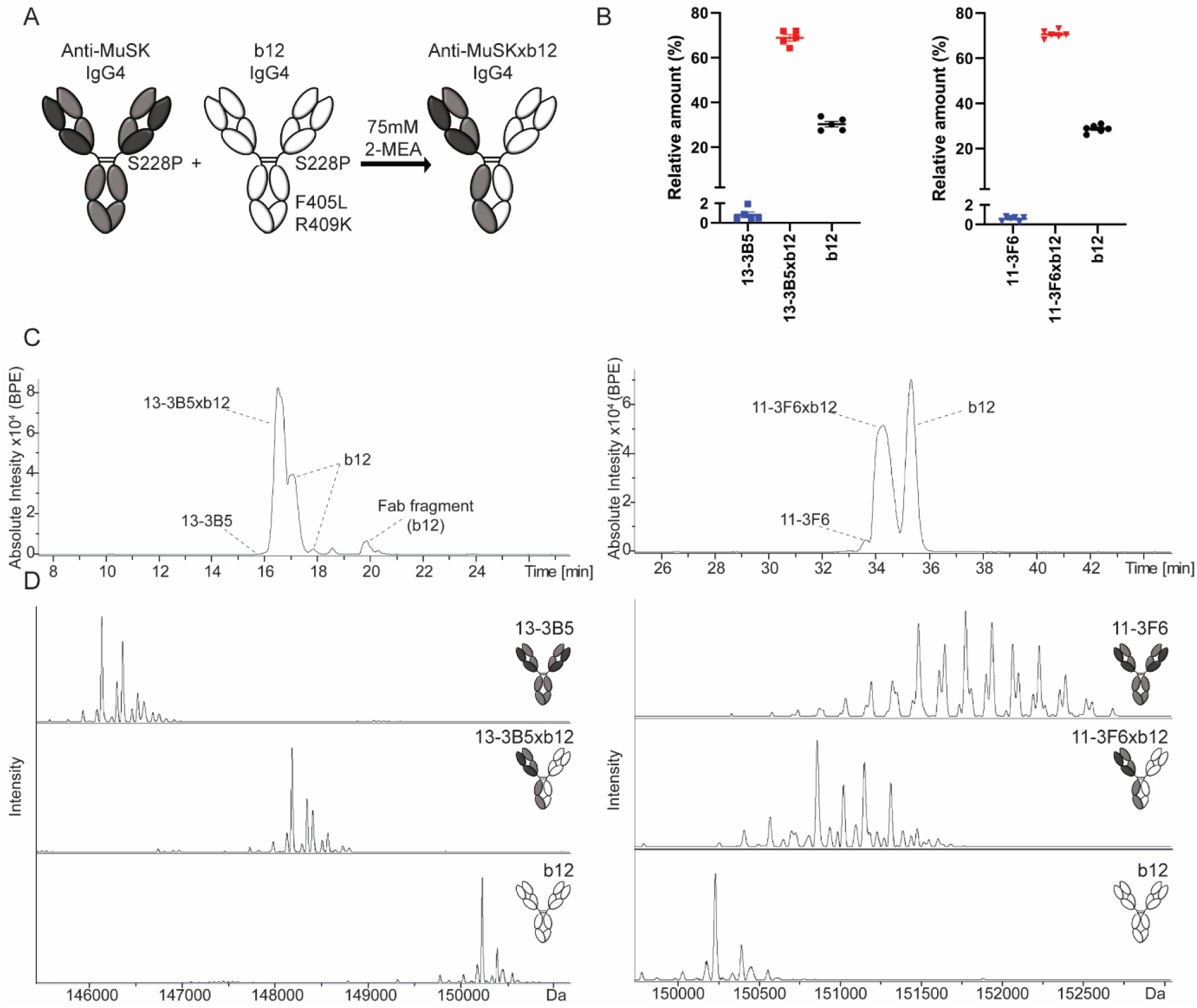
Generation of stable, MuSK MG patient-derived, bispecific, monovalent recombinant IgG4 MuSK antibodies. (A) Graphic depiction of the amino acid substitutions introduced for controlled Fab-arm exchange with IgG4. To make all antibodies resistant to Fab-arm exchange *in vivo*, serine 228 was converted to a proline. To stabilize bispecific (monovalent) IgG4s a leucine (L) at position 405 and lysine (K) at position 409 were introduced in the b12 antibody to create complementary amino acids with the phenylalanine (F) and arginine (R) in the anti-MuSK IgG4s respectively. (B) The exchange reaction with 75 mM 2-mercaptoethylamine (2-MEA) yielded high exchange efficiency with <1% contamination of the original monospecific (bivalent) anti-MuSK IgG4 for both clones (n=5). The b12 exchange partner was added in excess, yielding a remnant of ~30% after exchange. Representative examples of (C) base peak electropherograms (BPE) and (D) deconvoluted mass spectra of 11-3F6 and 13-3B5 respectively. The materials for 11-3F6xb12 exchange efficiency tests came from productions in both CHO and HEK293 cells. Representative images of 11-3F6 and 11-3F6xb12 deconvoluted mass spectra are from a production in CHO cells. Data represents mean ± SEM.

The exchange efficiency and residual amount of the bivalent (monospecific) anti-MuSK antibodies were determined with capillary electrophoresis (CE) hyphenated with mass spectrometry (MS) (Gstöttner et al., 2020). The monovalent (bispecific) IgG4 was separated from its two bivalent parents with CE, permitting reliable determination of their relative amounts down to 0.5% (Fig. 1C, Supplementary Fig. 1). The purity of the separated fractions was confirmed with MS (Fig. 1D). The efficiency of exchange was high for both antibodies, with <1% of remaining bivalent MuSK antibody. The b12 antibody was added in molar excess to drive exchange of the MuSK antibody to completion and on average ~30% excess b12 IgG4 is therefore apparent in the CE and MS analyses (Fig. 1B).

### Monovalent MuSK antibodies inhibit MuSK signaling in vitro

C2C12 myotubes contain all the muscle-specific machinery to interrogate the agrin-Lrp4-MuSK-signaling cascade *in vitro*, except for neural agrin. Addition of agrin to these cultures activates this cascade, leading to MuSK phosphorylation and AChR clustering (Fuhrer et al., 1997; Glass et al., 1996; Zhou et al., 1999). Monovalent MuSK antibodies were found to inhibit agrin-induced MuSK phosphorylation in a concentration-dependent manner (Fig. 2, A and B). Bivalent anti-MuSK IgG4 in stark contrast (partially) induced MuSK phosphorylation and AChR clustering both in the absence or presence of agrin (Fig. 2, C-F). These opposing effects extend previous observations using monovalent Fab fragments (Fichtner et al., 2020; Huijbers et al., 2019b). The bivalent 13-3B5 and 11-3F6 antibodies induced MuSK phosphorylation to different extents, with 13-3B5 reaching supra-agrin levels, while maximum phosphorylation induced by 11-3F6 reached about 70% of the level of agrin. Bivalent anti-MuSK antibodies induced similar amounts of large AChR clusters (>15 μm^2^), which are considered most mature and therefore relevant (Phillips et al., 2015). However, 13-3B5 also induced more smaller clusters compared to 11-3F6 (Supplementary Fig. 2A). In summary, monovalent MuSK IgG4 abolished agrin-Lrp4-MuSK signaling *in vitro*, whereas bivalent anti-MuSK IgG4 partially activated this signaling, independent of agrin.

**Fig. 2.**
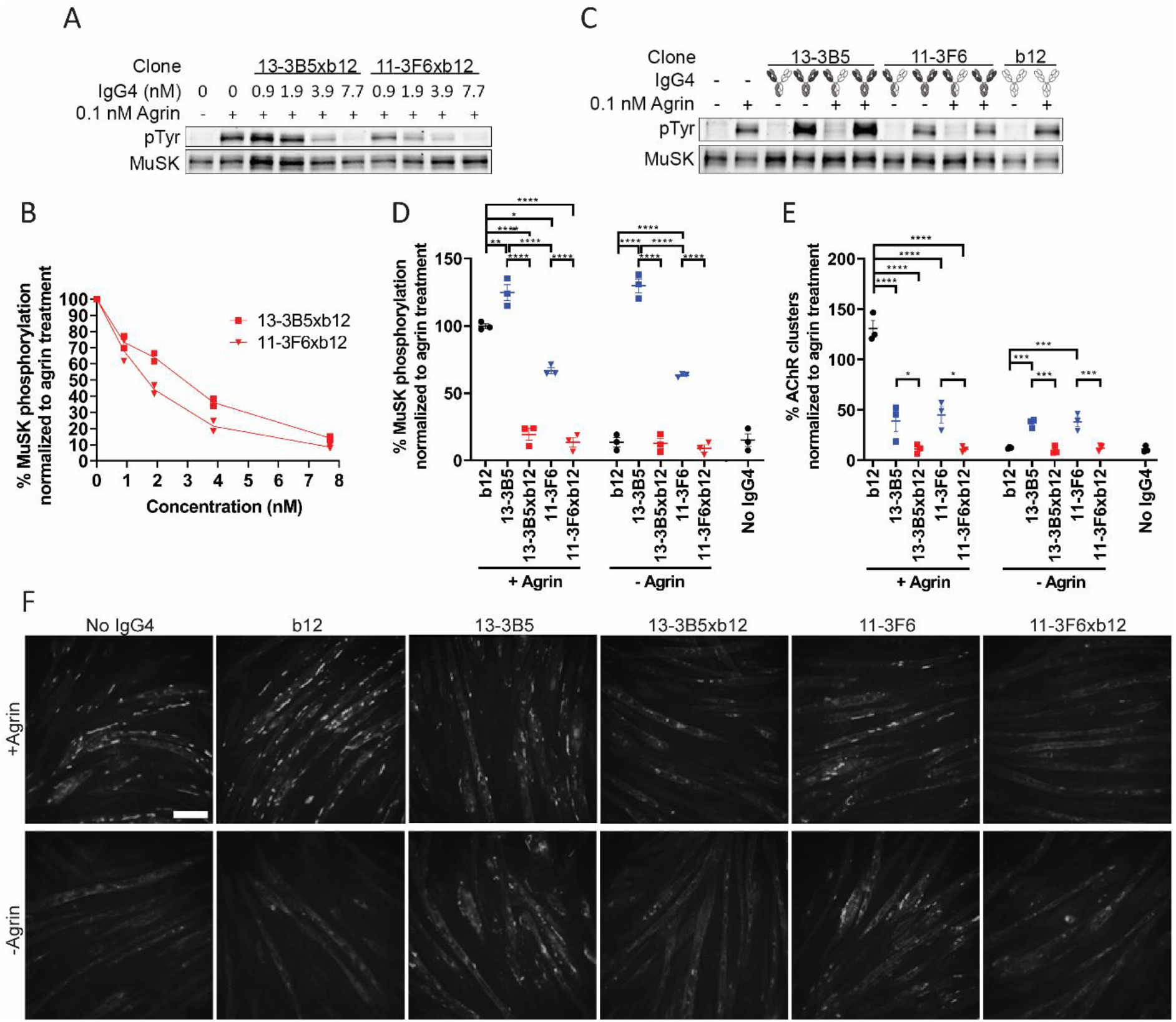
Monovalent IgG4 MuSK antibodies abolish agrin-induced signaling of the agrin-Lrp4-MuSK cascade. (A and B) Monovalent MuSK antibodies inhibited agrin-induced MuSK phosphorylation in a concentration-dependent manner in C2C12 myotubes (n=2). Maximum inhibition was reached at 7.7 nM. (C and D) Bivalent MuSK antibodies activated MuSK phosphorylation independent of agrin, while monovalent MuSK antibodies fully inhibited agrin-induced MuSK phosphorylation. The b12 control did not affect MuSK phosphorylation. To correct for loading differences the phosphotyrosine (pTyr) signal was divided by the amount of MuSK that was immunoprecipitated and normalized to the agrin only condition per replicate (n=3). (E and F) Monovalent MuSK antibodies completely inhibited agrin-induced AChR clustering (visualized by AF488-BTX staining). Bivalent MuSK antibodies partially induced AChR clustering independent of agrin, but partially inhibited agrin-induced clustering. Large (>15 μm^2^) AChR clusters were counted and normalized to the agrin only condition per replicate (n=3). Data represents mean ± SEM. One-way ANOVA with Šidák-corrected comparisons (D and E) * <0.05, ** <0.01, *** <0.001, **** <0.0001. Scalebar=100 μm.

### Monovalent MuSK antibodies are more pathogenic than bivalent MuSK antibodies in mice

Passive transfer of MuSK antibodies to NOD/SCID mice is a well-established method to investigate the development of myasthenic muscle weakness without confounding immune reactions to human antibodies (Huijbers et al., 2019a; Klooster et al., 2012). Immunostaining of whole-mount levator auris longus muscle confirmed that both mono- and bivalent MuSK antibodies bound MuSK at the NMJ, while the control b12 antibody did not (Supplementary Fig. 2B). The *in vivo* half-life ranged between 38-63 hours and varied by antibody (Supplementary Fig. 3, A and B). To ensure continuous *in vivo* exposure, an injection regimen of every 3-4 days was chosen for the passive transfer experiments. The minimum dose required to induce progressive phenotypical myasthenic symptoms was determined with monovalent 11-3F6xb12 IgG4. A dose of 2.5 mg/kg every 3-4 days resulted in progressive muscle weakness starting after about 5 days, leading to 20% body weight loss after 11 days (Supplementary Fig. 3, D-F). No clinical myasthenic muscle weakness or weight loss was observed at the 1.25 mg/kg dose. The serum antibody levels in these latter mice were approximately ten-fold lower compared to the 2.5 mg/kg regimen, indicating antigen-mediated disposition at lower doses (Supplementary Fig. 3C). To compare the pathogenicity of monovalent vs. bivalent MuSK antibodies, we injected the bivalent or monovalent MuSK antibodies, or the b12 control at 2.5 mg/kg every three days (Fig. 3A).

**Fig. 3.**
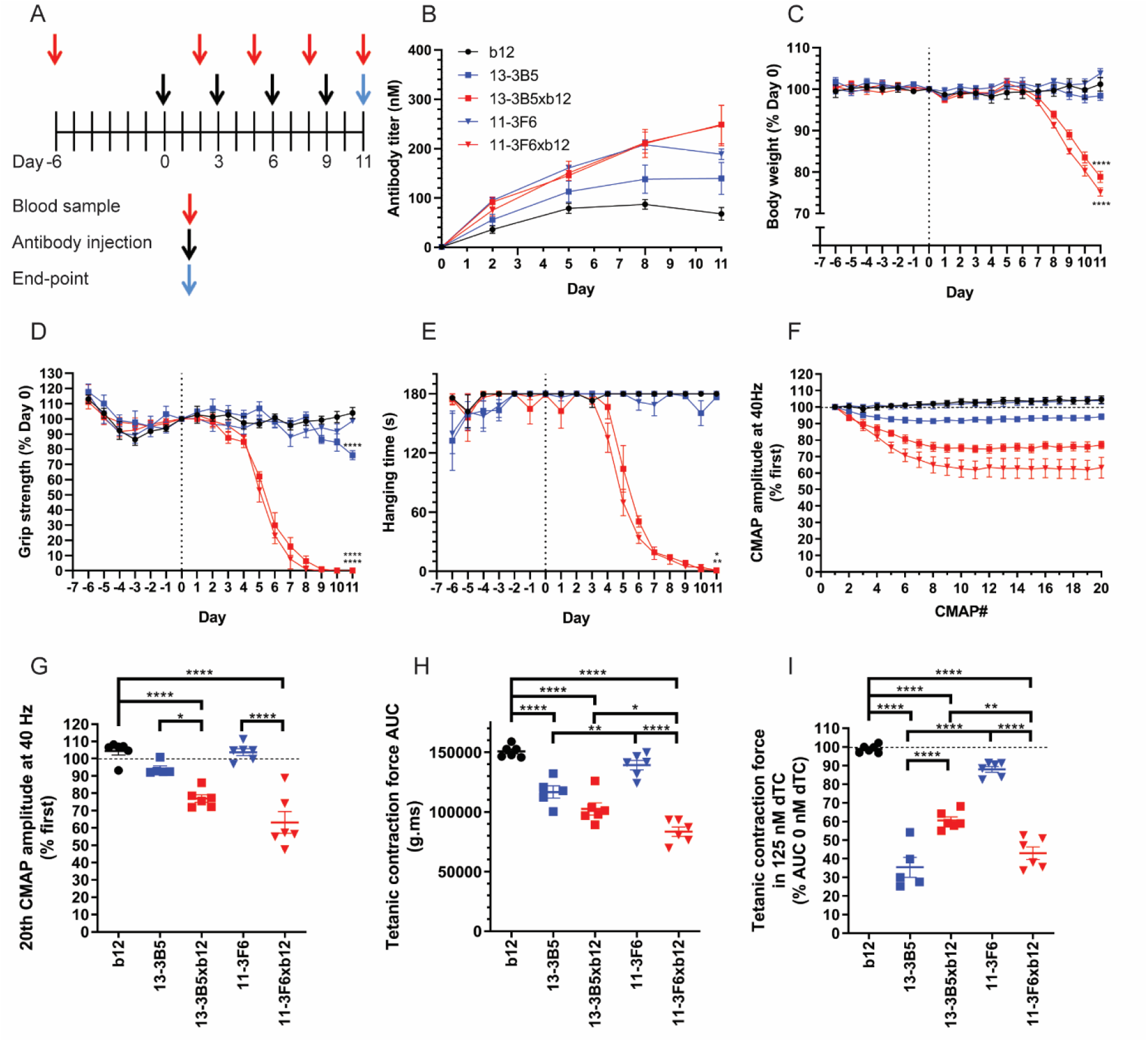
Monovalent MuSK antibodies induce rapid onset, progressive myasthenic symptoms in mice, whereas bivalent MuSK antibodies do not. (A) Experimental design of passive transfer with 2.5 mg/kg recombinant antibody (B) Serum antibody titers confirmed exposure. (C-E) Monovalent, but not bivalent, MuSK antibodies induced progressive weight loss and muscle weakness. 13-3B5 induced a delayed, mild loss of grip strength. (F and G) Monovalent, but not bivalent, MuSK antibodies induced a significantly larger CMAP decrement upon repetitive stimulation at 40 Hz compared to the b12 control. (H) *Ex vivo* tetanic contraction force of the diaphragm was reduced by 13-3B5, 13-3B5xb12 and 11-3F6xb12 compared to b12. (I) The safety factor of neuromuscular transmission was assessed in the presence of 125 nM dTC. Exposure to all MuSK antibodies reduced the safety factor compared to the b12 control, although for 11-3F6 this was only a trend (p=0.079). 11-3F6 and 11-3F6xb12 n=6, 13-3B5 n=5, 13-3B5xb12 and b12 n=6 (hanging time n=5). Data represents mean ± SEM. One-way ANOVA with Šidák-corrected comparisons for all parameters (C, D, G, H and I) except hanging time, for which Kruskal-Wallis test was used (C). * <0.05, ** <0.01, *** <0.001, **** <0.0001 compared to b12 group, unless otherwise specified.

Both monovalent anti-MuSK IgG4s induced rapid and severe myasthenic muscle weakness and body weight loss (Fig. 3, C-E and Supplementary Fig. 4, B-D for individual traces). The mice lost nearly all their grip strength and showed progressive fatigable muscle weakness in the inverted mesh test, starting within one week (Fig. 3, D and E). After a week, the mice also started to rapidly lose weight, likely as a consequence of the previously described bulbar muscle weakness, which makes it difficult to chew and swallow food (Fig. 3C) (Evoli et al., 2003; Morsch et al., 2012; Zhou et al., 2004). In sharp contrast, the bivalent 11-3F6 and 13-3B5 did not induce weight loss. Bivalent 11-3F6 furthermore did not induce any sign of clinical muscle weakness in grip strength or inverted mesh hanging time (Fig. 3, D and E). The bivalent 13-3B5 IgG4 induced a mild, but statistically significant, loss of grip strength only at the final day of the experiment (Fig. 3D). Furthermore, three out of five mice in this group could not complete the three minutes hanging test on day 10 and/or 11, while all mice in the control group could (Fig. 3E and Supplementary Fig. 4D).

To assess muscle function in more detail, repetitive nerve stimulation electromyography (EMG) on the left calf muscles was performed on the final day of the experiment. Decrement of the compound muscle action potential (CMAP) is a clinical electrophysiological hallmark of MG, and is caused by loss of transmission in a progressive number of NMJs during the stimulations (Plomp et al., 2015). Both monovalent MuSK antibodies induced 20-30% decrement at 40 Hz simulation (approximately the physiological firing rate of motor neurons (Eken, 1998)), indicating severe muscle fatigability (Fig. 3, F and G). This decrement was significantly larger compared to the bivalent parent of both IgG4s. Notably, the bivalent antibodies did not differ from the b12 control group, indicating normal muscle functioning.

To monitor *in vivo* exposure to the antibodies, serum samples were taken before the first injection and two days after every following injection. Serum titers varied between individual mice (Fig. 3B and Supplementary Fig. 4A), indicating individual differences in pharmacokinetics. Specifically, serum levels of the b12 antibody were lower compared to the different anti-MuSK clones, in spite of identical dosing. The levels of the MuSK antibodies were quite comparable, except at the final stage of the experiment, where titers of the bivalent IgG4s plateaued, while the monovalent anti-MuSK IgG4 serum levels continued to rise. This was probably due to the body weight and volume loss which mice injected with monovalent MuSK antibodies experienced at this stage.

To confirm that the IgG4s were stable *in vivo*, serum was collected at the final day of the experiment and analyzed with CE-MS. All five antibodies were successfully retrieved from the circulation and found to be intact at the end of the experiment (Supplementary Fig. 5). The only modification observed was the *in vivo* removal of C-terminal lysine for 11-3F6 and 11-3F6xb12 (Supplementary Fig. 5, C and E). C-terminal lysine clipping of IgGs is a well-known natural process, which does not affect antigen-binding ability (Liu et al., 2014). For both monovalent MuSK antibodies the proportion of excess b12 antibody seems to be reduced after injection, as seen in the base peak electropherogram (BPE) (Supplementary Fig. 5, D and E). The parental b12 control antibody therefore seems to be cleared slightly faster, but this is unlikely to have had an effect on the model as such. No other species or degradation products could be detected, indicating that the antibodies remained stable *in vivo* and ruling out confounding effects, like disassembly of bivalent MuSK antibodies.

In conclusion, all mice exposed to monovalent MuSK IgG4 rapidly developed severe myasthenic symptoms on all *in vivo* outcome measures. Mice exposed to the same dosing of bivalent MuSK IgG4 did not show overt phenotypical myasthenia. Thus, the functional monovalency of these bispecific (Fab-arm exchanged) anti-MuSK IgG4s makes them much more pathogenic than their monospecific equivalents with functional bivalency for MuSK.

### Monovalent MuSK antibodies induce reduced ex vivo contraction force compared to bivalent MuSK antibodies

To further characterize muscle function, the contraction force upon repetitive nerve stimulation was examined on the left hemidiaphragm *ex vivo*. Stimulation of the phrenic nerve at 40 Hz for 7 s resulted in tetanic contraction. Quantification of the area under the curve (AUC) revealed similar patterns between groups as seen with the CMAP decrement in the calf muscles. The monovalent MuSK antibodies reduced the contraction force most severely (Fig. 3H). Baseline contraction force upon exposure to bivalent 11-3F6 was indistinguishable from the b12 control group; while mice that received bivalent 13-3B5 showed a mild but statistically significant reduction of contraction force. Furthermore, the diaphragm contraction force was lower for the monovalent compared to their bivalent counterparts; although this was only statistically significant for 11-3F6xb12.

Subclinical signs of myasthenic muscle pathophysiology can be investigated by assessing the safety factor of neuromuscular transmission with the reversible AChR blocker d-tubocurarine (dTC). In the b12 control group, the tetanic contraction force of the diaphragm was not reduced by 125 nM dTC, indicating a healthy safety factor (Fig. 3I). Both monovalent anti-MuSK IgG4s substantially reduced the safety factor. In contrast, bivalent 11-3F6 did not show a statistically significant reduction of tetanic contraction, although a trend was seen (p=0.079). Diaphragms of mice exposed to bivalent 13-3B5 were affected by 125 nM dTC, indicating that although the clinical phenotype caused by 13-3B5 is mild, the NMJs have already substantial loss of AChR and are in a state of subclinical myasthenia (Fig. 3I). Taken together, monovalent MuSK IgG4 antibodies caused significant reduction in contraction force and safety factor of neurotransmission. Bivalent MuSK antibodies induced subclinical signs of myasthenia, but the extent was antibody-dependent.

### Monovalent and bivalent MuSK antibodies differentially affect neuromuscular junction morphology

Fragmented and reduced AChR area at NMJs is a well-described pathological feature, causing the muscle weakness in MuSK MG animal models (Cole et al., 2008; Ghazanfari et al., 2014; Jha et al., 2006; Klooster et al., 2012; Mori et al., 2012a; Morsch et al., 2012; Punga et al., 2011; Richman et al., 2012). To investigate the postsynaptic NMJ morphology after *in vivo* exposure to mono- and bivalent MuSK antibodies, AChRs were stained in whole-mount preparations of the right hemidiaphragm and epitrochleoanconeus (ETA) (Fig. 4 and Supplementary Fig. 6A and D). Quantification of the total AChR signal per NMJ revealed that both monovalent anti-MuSK IgG4s and the bivalent 13-3B5 caused a strong reduction of AChRs, while mice exposed to bivalent 11-3F6 had more remaining AChRs compared to monovalent 11-3F6xb12 and bivalent 13-3B5 (Fig. 4A). The area of positive AChR signal was also significantly reduced in the muscles from mice treated with both monovalent anti-MuSK IgG4s and bivalent 13-3B5 (Fig. 4B). The average intensity of post-threshold AChR staining was not statistically different between the conditions (Fig. 4C). NMJs of mice exposed to both monovalent anti-MuSK IgG4s appear fainter, but the pretzel-like structure remained relatively intact (Fig. 4D). In contrast, the pretzel-like structure of NMJs appear fragmented upon exposure to bivalent 13-3B5. In sum, exposure to mono- and bivalent MuSK antibodies reduced the number of postsynaptic AChRs, with the effect of the bivalent MuSK antibodies being antibody-dependent.

**Fig. 4.**
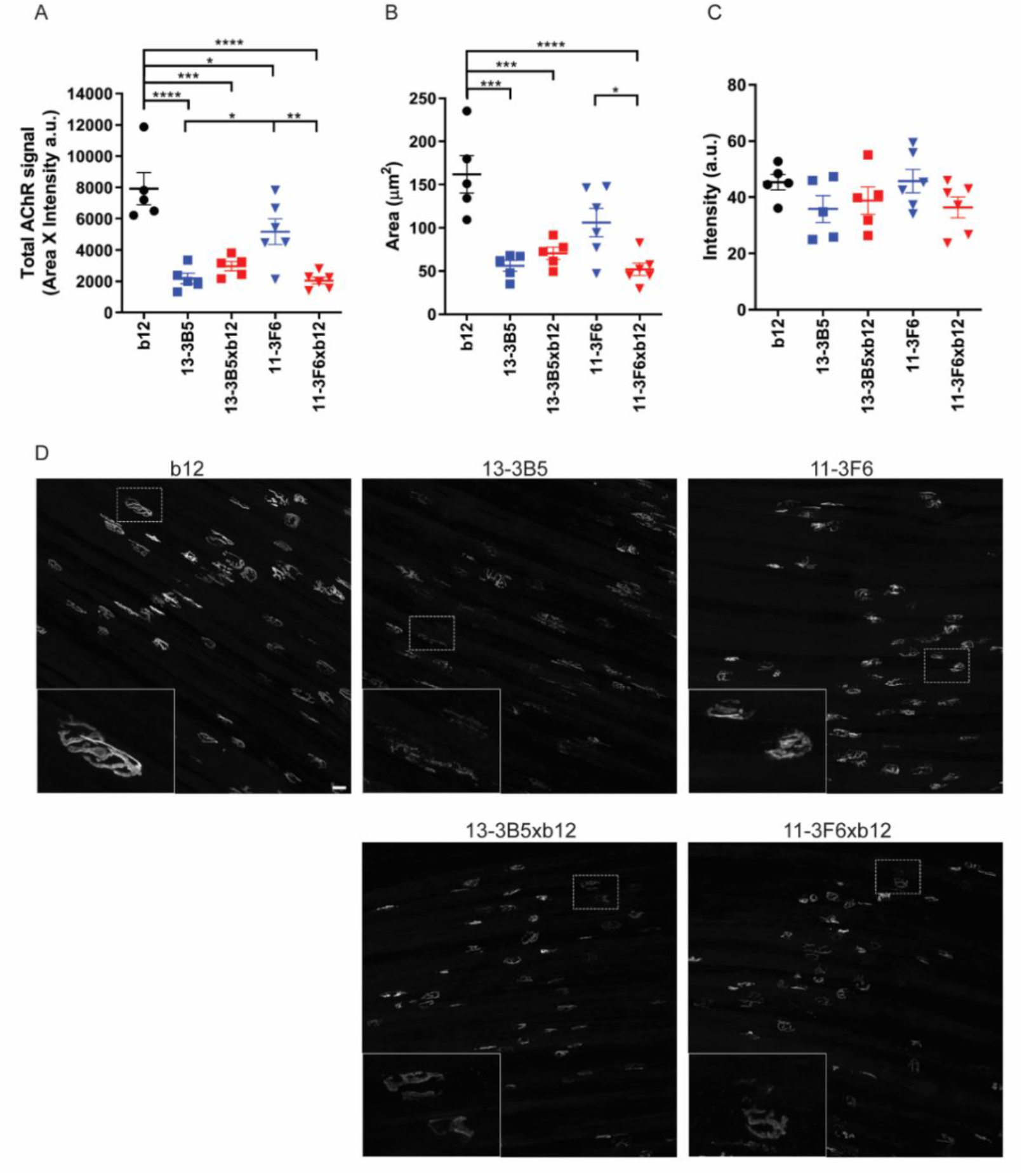
Monovalent and bivalent MuSK antibodies impair NMJ morphology to a different extent. To visualize the postsynaptic NMJ, AChRs were stained with AF488-BTX on diaphragm muscle preparations. Twenty randomly selected NMJs per diaphragm were analyzed and averaged. (**A**) Total AChR signal was calculated by multiplying the positive area with the average intensity per NMJ. Both bivalent and monovalent MuSK antibodies significantly reduced the total AChR signal compared to the b12-treated mice. Bivalent 13-3B5 and monovalent 11-3F6xb12 reduced the total AChR signal to a greater extend compared to monospecific 11-3F6. Bivalent 13-3B5 did not significantly differ from monovalent 13-3B5xb12. (**B**) To assess the size of the NMJs, a threshold was applied and the area of the positive signal was quantified. Exposure to both monovalent MuSK antibodies and bivalent 13-3B5 resulted in less signal reaching the threshold and therefore smaller NMJs. Monovalent 11-3F6xb12 also caused significantly smaller NMJs compared to bivalent 11-3F6. (**C**) The average intensity of AChR staining surpassing the threshold did not significantly differ between groups. (**D**) representative maximum projections per condition with insets. Scalebar = 25 μm. 11-3F6 and 11-3F6xb12 n=6, 13-3B5, 13-3B5xb12 and b12 n=5. Data represents mean ± SEM. One-way ANOVA with Šidák-corrected comparisons for all parameter * <0.05, ** <0.01, *** <0.001, **** <0.0001.

Loss of appropriate pre- and postsynaptic alignment is also known to contribute to the NMJ dysfunction seen in MG animal models (Cole et al., 2008; Huijbers et al., 2018; Klooster et al., 2012). Therefore, this was assessed in the ETA muscle of a subset of animals (Supplementary Fig. 6). The intensity and morphology of the presynaptic SV2 signal were not affected by exposure to these mono- and bivalent anti-MuSK IgG4s (Supplementary Fig. 6B). Denervation assessed by the colocalization of presynaptic SV2 signal with postsynaptic AChR signal revealed a similar pattern as seen with the postsynaptic AChR signal (Supplementary Fig. 6 A and C). Overall, this data suggests that postsynaptic pathology of the NMJ is the main cause of the neuromuscular dysfunction induced by these MuSK antibodies and that loss of AChR area results in less alignment with the pre-synapse.

### Antibody-dependent pathogenicity of parental bivalent MuSK antibodies

To investigate whether both bivalent MuSK antibodies could be pathogenic, albeit at a higher dose or with prolonged exposure, we increased the dosing to 5 mg/kg and 10 mg/kg, and extended the duration of the passive transfer to 3 weeks (Fig. 5A). The mice exposed to bivalent 13-3B5 displayed overt signs of muscle weakness on grip strength and inverted mesh from day 9-12 and progressively lost weight in week three (Fig. 5, B-D and Supplementary Fig 7, B-D for individual traces). Grip strength slowly declined over a period of a week to a near complete loss on the final days of the experiment (Fig. 5C). Hanging time on the inverted mesh did not show a clear progressive decline in the majority of mice exposed to bivalent 13-3B5 (Supplementary Fig. 7D). Lastly, exposure to bivalent 13-3B5 resulted in a significant CMAP decrement compared to both the control group and bivalent 11-3F6 (Fig. 5E). In sum, prolonged exposure to bivalent 13-3B5 IgG4 can cause progressive and severe myasthenic muscle weakness. Interestingly, the onset of symptoms is later and progresses slower compared to monovalent 13-3B5xb12.

**Fig. 5.**
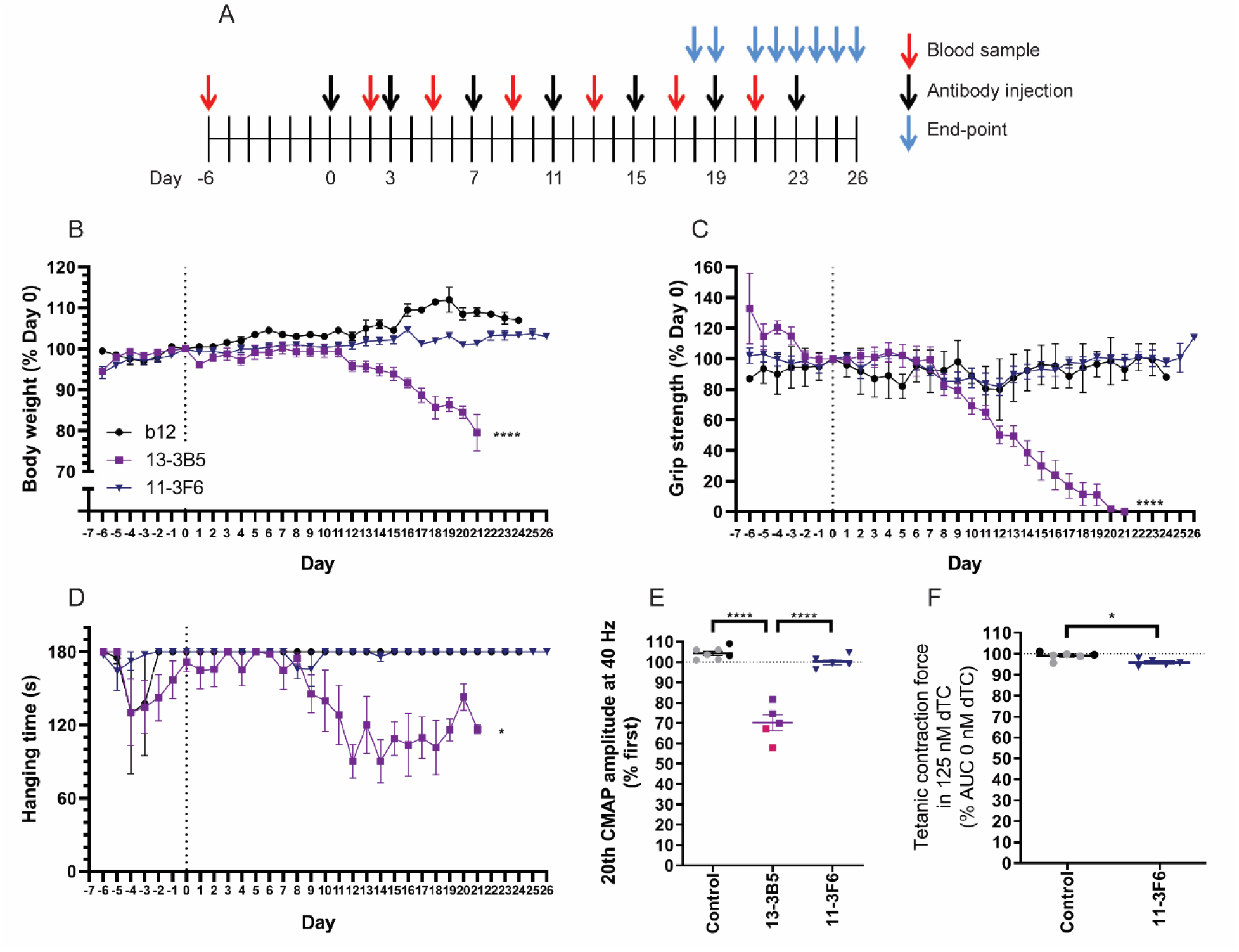
Antibody-dependent pathogenicity of bivalent MuSK antibodies. (**A**) Experimental design of passive transfer in NOD/SCID mice. Mice exposed to bivalent 13-3B5 started to (progressively) lose weight (**B**), grip strength (**C**) and hanging time on the inverted mesh (**D**) in the third and second week of the experiment respectively. Mice exposed to bivalent 11-3F6 did not show progressive loss on these parameters (**B-D**). (**E**) Mice exposed to bivalent 13-3B5 showed a significant ~30% CMAP decrement at the endpoint, while bivalent 11-3F6 did not induce a decrement. (**F**) Contraction force of the diaphragm was significantly, but very mildly (~3% reduction) affected by 125 nM dTC for mice exposed to bivalent 11-3F6. Data represents mean ± SEM. 13-3B5: 5 mg/kg (pink) n=2, 10 mg/kg (purple) n=4 (except for hanging time on inverted mesh and EMG n=3); 11-3F6: 10 mg/kg n=5; b12 (black): n=2, combined with untreated (grey) n=5 for CMAP or n=4 for tetanic contraction).

Mice exposed to bivalent 11-3F6 did not show signs of myasthenic muscle weakness on any of the *in vivo* parameters at higher dose and after prolonged exposure (Fig. 5B-E). To detect possible subclinical pathology, the safety factor of neurotransmission was assessed in the diaphragm. For reference, the relative tetanic contraction force of animals treated with the b12 antibody was supplemented with untreated healthy animals. The safety factor of mice exposed to 10 mg/kg bivalent 11-3F6 slightly (~3%), but significantly reduced compared to control muscle (Fig. 5F). The effect size of the reduced safety factor seen in mice exposed to 2.5 mg/kg bivalent 11-3F6 for 11 days therefore does not seem to worsen over time and with higher dose (Fig. 3I). Antibody titers were comparable between mice injected with bivalent 13-3B5 or 11-3F6 (Supplementary Fig. 7A). The differential effects between 13-3B5 and 11-3F6 therefore confirms bivalent MuSK antibodies can be pathogenic, but their pathogenic potential is antibody-dependent.

## Discussion

In this study, we demonstrate a new disease mechanism in autoimmunity related to the unique feature of IgG4 to undergo Fab-arm exchange. We provide evidence that autoantibody functional monovalency for MuSK amplifies the *in vivo* pathogenicity of IgG4 MuSK antibodies. Mechanistically, this may be explained by the respective antagonistic vs. agonistic effects of the Fab-arm exchanged monovalent and parental bivalent IgG4 antibodies on the MuSK signalling cascade. Thus, class switching to IgG4 may not just be a characteristic of IgG4-mediated autoimmunity, but also be a crucial step in symptom manifestation.

IgG4 is the only human antibody subclass able to undergo Fab-arm exchange under physiological conditions (Rispens et al., 2014). It furthermore has limited ability to stimulate inflammation. The relevance of (Fab-arm exchanged) IgG4 is thought to be inhibition of ongoing detrimental immune responses against exogenous antigens and allergens (Lighaam and Rispens, 2016). For example, in novice bee-keepers prolonged exposure to bee venom induces isotype/subclass switching to IgG4 which dampens allergic responses, rendering beekeepers resistant to bee stings (Aalberse et al., 1983). Moreover, chronic inflammation due to worm infections is halted by class switching to IgG4 (Adjobimey and Hoerauf, 2010). Why certain (auto)immune responses undergo a class switch to predominant IgG4 responses is not known. In MuSK MG, autoantibodies of the IgG1, 2 or 3 subclasses are monospecific and may activate complement. However, given their significantly lower levels in serum and potential agonistic effects, their contribution to the clinical manifestation in patients is uncertain (McConville et al., 2004). It is possible that in IgG4-mediated autoimmune diseases a mild IgG1-IgG3 immune response against the antigen is ongoing, similar to bee keepers. This prolonged exposure at some point may induce an autoantibody class switch to high titers of functionally monovalent IgG4 and only then symptoms manifest. In the development of MuSK MG, high affinity binding of MuSK antibodies may furthermore be extra critical, as functional monovalent (germlined) monoclonal MuSK antibodies lost significant binding capacity and *in vitro* pathogenicity (Fichtner et al., 2020). Affinity maturation thus seems an additional requirement for potentiating pathogenicity of monovalent MuSK antibodies. It will be exciting to learn what governs the development of IgG4 (auto)immunity.

Monovalent anti-MuSK IgG4s inhibited agrin-Lrp4-MuSK signaling *in vitro* and induced progressive myasthenic weakness and NMJ morphological abnormalities in NOD/SCID mice similar to polyclonal patient-purified IgG4 and monovalent Fab fragments (Fichtner et al., 2020; Huijbers et al., 2019b; Huijbers et al., 2013; Klooster et al., 2012; Koneczny et al., 2013; Mori et al., 2012b). This suggests that Fab-arm exchanged anti-MuSK IgG4 in polyclonal patient IgG is the main pathogenic factor causing muscle weakness in patients and animal models. Importantly, this is the first study using patient-derived monoclonal monovalent antibodies to model MuSK MG *in vivo*. These models are exciting new tools to perform preclinical therapeutic tests.

Patient-derived bivalent MuSK antibodies can potentially be pathogenic *in vivo*, but this differed considerably between the two antibodies studied here. The bivalent parent antibody 13-3B5 was able to cause overt myasthenic symptoms, but this required more time to start (2-3 weeks) and progressed more slowly, as compared to monovalent 13-3B5xb12. This time-course of myasthenic symptom development and progression resembles what has been reported in MuSK active immunization models (Mori et al., 2012a; Patel et al., 2014; Punga et al., 2011; Ulusoy et al., 2014; Viegas et al., 2012). The MuSK antibodies induced during active immunization are functionally bivalent and monospecific, as rodent IgG is unable to exchange Fab-arms under physiological conditions (Lighaam and Rispens, 2016). Furthermore, MuSK immunization of mice induces a dominant mouse IgG1 response (Ulusoy et al., 2014). Mouse IgG1 activates complement weakly (Neuberger and Rajewsky, 1981), suggesting that the effects of direct binding are the main cause of myasthenic symptoms upon active immunization. Therefore, the mechanism of MuSK MG in active immunization models is expected to resemble that of pathogenic monospecific, functionally bivalent MuSK antibodies like 13-3B5. Taking together the opposing effects on MuSK signaling *in vitro* and the different timeline and development of muscle weakness and postsynaptic NMJ pathology *in vivo*, it would be interesting to investigate whether bivalent 13-3B5 is pathogenic through a different mechanism compared to the monovalent anti-MuSK IgG4s.

The bivalent 11-3F6 parent antibody did not induce any signs of clinical myasthenic muscle weakness, not even when doubling the exposure time and increasing the dose. Subtly different epitopes may differentially affect the conformation and activation of MuSK, and thereby the subsequent downstream signaling. Indeed 13-3B5 and 11-3F6 bind different epitopes (data not shown). These observations indicate that epitope specificity and antigen-manipulation is crucial for the pathogenicity of bivalent MuSK antibodies. Whether non-pathogenic MuSK antibodies such as 11-3F6 or #13 (Xie et al., 1997), can mitigate detrimental effects of pathogenic mono- and bivalent MuSK antibodies is not known.

It is important to note that in patients a polyclonal pool of MuSK antibodies exists. Some of these may be functionally bivalent, for example when MuSK antibodies are of the IgG1, IgG2 or IgG3 subclasses or non-exchanged IgG4. These antibodies may be non-pathogenic (like 11-3F6) and may compete for binding with inhibitory monovalent anti-MuSK IgG4. Alternatively, some of these may be pathogenic (like 13-3B5) and together with monovalent anti-MuSK IgG4 induce disease. The net result on NMJ function and disease severity will depend on the complex combination of these antagonistic and agonistic effects. It is further important to realize that the (lack of) pathogenicity of the bivalent human MuSK antibodies in this study can be only mediated by a direct effect on MuSK, as the model system used does not allow for assessment of for example complement activation (Schultz et al., 1995). This means it is still possible that if the subclass of the bivalent antibody allows for complement activation, it might be pathogenic in other model systems or in humans.

A critical step in this study was the generation of stable bispecific (monovalent) IgG4 antibodies. The cFAE method presented here broadens the toolbox available for generating therapeutic bispecific antibodies. By altering three residues in the Fc tail of IgG4 and adding excess of an irrelevant donor antibody (b12), <1% monospecific antibody contamination could be achieved. Furthermore, these bispecific IgG4s were stable after intraperitoneal injection in NOD/SCID mice. The field of bispecific antibody therapeutics has taken an exponential flight (Labrijn et al., 2019). The therapeutic promise of bispecific antibodies lies in their ability to bring together two antigens that could not be brought in close proximity with monospecific bivalent antibodies. Because of their naturally flexible structure and anti-inflammatory nature, stable bispecific IgG4 have shown promise in preclinical tests for T cell redirection in the treatment of cancer and hemophilia A (Ishiguro et al., 2017; Nair-Gupta et al., 2020; Smith et al., 2016). Our method adds to other currently available (IgG4) antibody technology platforms and can be used for development of antibody therapies in the future.

Taken together, IgG4 Fab-arm exchange is not an innocent bystander activity, but instead potentiates the pathogenicity of IgG4 autoantibodies. Our data show that rendering the IgG4s bispecific and thereby functionally monovalent for the MuSK autoantigen serves as a mechanism that exacerbates the autoantibodies pathogenic potential. This may be relevant for the growing number of IgG4-mediated autoimmune diseases and other disease settings where IgG4 plays a pathogenic role, such as where IgG4 blocks endogenous anti-tumor responses in melanoma patients (Daveau et al., 1977; Huijbers et al., 2018; Karagiannis et al., 2013).

## Methods

### Generation of bivalent and monovalent recombinant antibodies

Anti-MuSK clones 11-3F6 and 13-3B5 were previously isolated from a MuSK MG patient and produced with a human IgG4 Fc (Huijbers et al., 2019b). The S228P amino acid change in the anti-MuSK antibodies was achieved by converting AGC>CCC through side-directed mutagenesis based on the QuikChange II system (Agilent). To make the b12 antibody suitable as an exchange partner for cFAE, the sequence was modified to achieve S228P, F405L and R409K amino acid changes in an pcDNA3.1 IgG4 backbone. Heavy and light chain sequences of the b12 antibody were ordered at GeneArt (Thermo Fisher). All sequences were verified using Sanger sequencing.

Recombinant monoclonal antibodies were produced in suspension FreeStyle HEK293-F cells as described previously (Huijbers et al., 2019b) or using transient CHO-based expression (Evitria). IgG was purified with a HiTrap MabSelect SuRe protein A affinity column (GE Healthcare) on an AKTA Pure (GE Healthcare). Antibodies were dialyzed to PBS in dialysis cassettes (Thermo Fisher) or desalted to PBS using an HiPrep 26/10 column (GE Healthcare) on an AKTA Pure, filter sterilized and stored at −20°C until use. Recombinant antibodies 13-3B5 and b12 used in all experiments were produced in HEK cells. For 11-3F6, a batch produced in HEK cells was used in Figure 1 and 2 and supplemental Figure 1, 2 and 3. A batch of 11-3F6 produced in CHO cells was used in Figure 1, 3, 4 and 5, and supplemental Figure 4, 5, 6 and 7.

To generate monovalent MuSK antibodies, each of the anti-MuSK clones was combined with 1.3-1.4x molar excess of the b12 antibody and 75 mM 2-Mercaptoethylamine-HCl (Sigma-Aldrich) for five hours at 31°C (Labrijn et al., 2014). Preparations were dialyzed back to PBS using dialyzer cassettes, filter sterilized and stored at 4°C until use. Exchange efficiency and monovalent antibody purity were assessed using CE-MS, described below.

Concentrations of all recombinant antibodies were determined with nanodrop (ND-1000, v.3.8.1) using specific extinction coefficients predicted with the ProtPI tool based on the amino acid sequences (Supplementary Table 1). Furthermore, antigen-binding capacity was assessed using MuSK or gp120 enzyme-linked immunosorbent assay (ELISA), described below. Antibody integrity was confirmed using PageBlue Coomassie stain, according to the manufacturer instructions (Thermo Fisher).

### Capillary electrophoresis mass spectrometry

Sheathless CE-MS was employed to assess exchange efficiency and purity of monovalent antibodies. Analyses were carried out on a CESI 8000 instrument (Sciex) coupled to an Impact Qtof mass spectrometer (Bruker Daltonics) equipped with a nanoelectrospray source. Porous-tip capillaries (91 cm x 30 μm ID) were obtained from Sciex. Capillaries were coated using polyethylenimine (Gelest) following the protocol described by Sciex (Santos et al., 2015). The background electrolyte (BGE) consisted of 10% acetic acid or 30% acetic acid and 10% MeOH for 13-3B5(xb12) and 11-3F6(xb12), respectively. Before each run, the capillary was flushed for 4 min at 100 psi with the BGE. Separation was performed by applying −20 kV at 20°C. Samples were buffer exchanged to the relevant BGE using 30 kDa MWCO filters (Vivaspin, 3 cycles of 10000xg at 4 °C) and hydrodynamically injected for 15 s using 2.5 psi. The mass spectrometer was operated in positive ionization mode using a capillary voltage of 1200 V, a drying gas temperature of 120°C and a drying gas flow rate of 1.2 L/min. An ISCID energy of 100 eV was employed to obtain proper declustering of the antibodies. Quadrupole ion and collision cell energy were 5.0 and 20.0 eV, respectively. MS control and data acquisition and analysis were performed using QTOF control and data analysis software (Bruker Daltonics). Molecular mass determinations were performed using the Maximum Entropy deconvolution algorithm of the data analysis software. A baseline subtraction of 0.7 points was applied to the deconvoluted mass spectra.

Relative amounts of the antibodies were determined by integrating the area under the peaks observed in the BPE. For 13-3B5, the areas were directly employed to determine the relative amounts of the antibodies. For 11-3F6, different ionization efficiency was observed compared to b12 as consequence of the Fab glycans. Therefore, the relative amounts were determined by adding known amounts of the antibodies and interpolating in the corresponding calibration lines.

### MuSK and gp120 ELISA

MuSK or gp120 ELISA was used to assess antigen reactivity or quantify the serum titers of the recombinant antibodies in the NOD.CB17-Prkdcscid/J (NOD/SCID) mice. To measure mono- or bivalent variants of 11-3F6 and 13-3B5, 3 μg/mL of the complete extracellular region of MuSK, produced as described previously (Huijbers et al., 2016), was coated on MaxiSorp plates (Thermo Fisher). For the b12 antibody, MaxiSorp plates were coated with 1 μg/mL HIV-GP120 protein (Sigma-Aldrich). Samples were diluted in an eight-point two-fold dilution series. Serum samples started at 125x dilution in block. The original batch of antibody that was injected served as a standard in duplicate; the first dilution started at 0.5-1 nM. Mouse anti-human IgG4 (Nordic clone N315, Nordic MUbio) and rabbit anti-mouse-AP (D0314, Dako) were used as secondary antibodies and conjugate respectively. Plates were developed with pNPP (VWR) and the reaction was stopped using sodium hydroxide. All samples were tested in duplicate and quantified in SoftMax pro (version 7.0.3, Molecular Devices).

### Immunostaining of mouse NMJs using monovalent antibodies

To determine binding capacity of recombinant mono- and bivalent MuSK antibodies to mouse MuSK, levator auris longus muscles of NOD/SCID mice were immunostained with 2 μg/mL recombinant antibody overnight at 4°C. The preparations were co-stained for synaptic regions with 2 μg/mL AlexaFluor594-conjugated α-bungarotoxin (BTX) (B13423, Thermo Fisher) and imaged as described previously (Huijbers et al., 2019b).

### MuSK phosphorylation and AChR clustering

C2C12 myoblasts were obtained from cell line services, tested for mycoplasma contamination and maintained for maximum 7 passages after thawing. MuSK phosphorylation and AChR clustering was assessed as described previously (Huijbers et al., 2019b). Briefly, for MuSK phosphorylation differentiated C2C12 myotubes were treated for 30 minutes. The concentration of monovalent MuSK antibodies was titered to achieve complete inhibition of agrin-induced MuSK phosphorylation. C2C12 myotubes were treated with this concentration (7.7 nM) of recombinant mono- or bivalent antibodies in the absence or presence of 0.1 nM neural agrin (R&D systems). MuSK was immunoprecipitated from whole lysate and detected on western blot. For AChR clustering, C2C12 myotubes were treated for 16 hours with 7.7 nM recombinant antibodies in 96-well plates. After treatment, cells were stained with 2 μg/mL BTX-488 (B13422, Thermo Fisher) and 2 μg/mL Hoechst 33342 (H1399, Thermo Fisher) for 30 minutes at 37°C before fixation with 4% paraformaldehyde (PFA). Twenty fields divided over 5 wells per condition were randomly selected in the brightfield channel on a Leica AF6000 microscope. AChR cluster count and size were analyzed using ImageJ 1.52n. A manual threshold was set for each independent replicate. Large (>15 μm^2^) and all (>3 μm^2^) clusters were analyzed separately. Both assays were performed in triplicate.

### Mouse passive transfer studies

NOD/SCID mice were used to avoid a mouse immune response to the injected human recombinant IgG4. Mice were bred in the LUMC or purchased from Charles River. They were housed in sterile, individually-ventilated cages and provided with sterile food and drinking water *ad libitum*. Female mice were aged 8-10 weeks at the start of the experiment, unless otherwise specified.

To determine antibody half-life, mice received a single 5 mg/kg intraperitoneal (i.p.) injection of one of the recombinant antibodies. Antibody titer was monitored in the serum obtained from blood drawn via tail vein cut. Because of animal ethical regulations, daily blood withdrawal was not allowed. Therefore, blood was drawn from mice per treatment group in an alternating fashion. Mouse 1 had blood samples (~50 μL) drawn 8, 72 and 144 hours after the injection using a tail cut. Mouse 2 had blood samples drawn 3 days before and 24, 96 and 168 hours after the injection. Samples were allowed to clot at room temperature, centrifuged at 10,000 RPM for 3 min and serum was stored at −20°C until analysis. Antibody titers were measured using the MuSK and gp120 ELISA as described above. A one-phase exponential decay function with the plateau constrained at zero was fitted in GraphPad Prism (version 8.1.1) per mice to calculate the half-life. Both male and female NOD/SCID mice were used in this experiment.

The experimenters were blinded for the injected antibodies throughout all experiments and analyses. To determine the minimum dose at which monovalent MuSK antibodies could be fully pathogenic, 11-3F6xb12 was i.p. injected on day 0, 3, 7, 11, 15 and 19 at doses of 1.25, 2.5 or 5 mg/kg (of the body weight at the beginning of the experiment). In addition to body weight, *in vivo* muscle strength and endurance were assessed on a daily basis using a grip strength meter and inverted mesh hang test as described previously (Klooster et al., 2012). To familiarize the mice with handling and the tests, the daily *in vivo* measurements were started 4-6 days prior to the first injection. Blood samples were taken on day -2, 5, 13 and on the final day of the experiment. Mice were killed if they lost >20% of their body weight compared to the first day of injection or when they reached the end of the experiment.

To compare pathogenicity of monovalent and bivalent MuSK antibodies, mice were i.p. injected with 2.5 mg/kg recombinant antibody every three days. A roughly equal distribution of body weight between treatment groups was ensured during blinding. Mice were allocated to a treatment group by a lab member not involved in the experiment. Serum samples were taken on day -6 and two days after every injection. Body weight, *in vivo* muscle strength and endurance were assessed daily. On day 11, mice were subjected to the endpoint analyses described below.

To further interrogate possible pathogenic effects of bivalent MuSK antibodies at higher doses over longer time, mice were injected with 5 mg/kg or 10 mg/kg on day 0, 3, 7, 11, 15, 19 and 23. Serum samples were taken prior to the first injection and two days after every injection. Mice that received 13-3B5 were sacrificed between day 18 and 21 as they lost weight. Mice that received 11-3F6 were sacrificed sequentially between day 21 and 26, due to time restrictions of the end-point analyses. For one mice that received 11-3F6, the first injection did not lead to systemic exposure as it could not be detected in the serum. Therefore, day 0 was moved to the day of the second injection for this animal. Mice were subjected to repetitive nerve stimulation EMG on the endpoint day. For the mice without signs of muscle weakness on the *in vivo* outcome measures, the diaphragm was prepared for measurement of *ex vivo* contraction force, described below. Untreated non-littermate NOD/SCID mice (males and females aged 2-4 months) were combined with the two animals that received the b12 control used as a healthy reference for EMG and contraction measurements.

Over all experiments, two mice had to be excluded for all parameters, because the serum titers revealed lower antibody levels on two time-points in the experiment, likely due to misplacement of i.p. injections. This concerned one animal injected with 2.5 mg/kg 13-3B5 and one animal injected with 5 mg/kg 11-3F6. For a further three mice (2.5 mg/kg b12, 2.5 mg/kg 13-3B5xb12 and 10 mg/kg 13-3B5), the measurements on the inverted mesh had to be excluded, because the animals did not complete 180 s hanging during the training period.

### Endpoint analyses

After the daily measurements on the endpoint day, mice were subjected to repetitive nerve stimulation EMG of the calf muscles under anesthesia, described in detail previously (Klooster et al., 2012). One of the mice treated with 10 mg/kg bivalent 13-3B5 died from the anesthesia before EMG could be conducted. Substantial CMAP decrement can be measured at 40 Hz stimulation in passive transfer MuSK MG models (Klooster et al., 2012). Upon completion of the EMG, blood was collected by cutting the tail without recovery from anesthesia. Immediately thereafter, mice were killed by CO_2_ inhalation. During dissection more blood was collected via vena cava puncture. The serum from the tail and vena cava was pooled and stored at −20°C until further analysis. The right hemidiaphragm and ETA were dissected for NMJ morphological analyses described below.

To measure *ex vivo* contraction force, the left phrenic nerve-hemidiaphragm was prepared as described previously (Huijbers et al., 2019a). Briefly, the hemidiaphragm was equilibrated in Ringer’s medium. The phrenic nerve was supramaximally stimulated at 40 Hz for 7 seconds every 5 minutes until the preparation gave stable contraction. The safety factor of neuromuscular transmission was assessed by incubating the preparation with 125 nM dTC (Sigma-Aldrich) and stimulating at 40 Hz for 7 seconds every 5 minutes until the preparation gave stable contraction.

### Neuromuscular junction morphology

The most dorsal strip of the right hemidiaphragm and the whole ETA were pinned up in Sylgard lined dishes and fixed in 1% PFA in PBS for 30 minutes. All incubations were done at RT, unless otherwise specified. To enable parallel processing of all muscles in an experiment, they were stored floating in 1% PFA at 4°C for 3-7 days. Before staining, muscles were blinded to the phenotypes observed in the experiment until all morphology analyses were completed. The tissues were extensively washed with PBS, remaining 1% PFA was neutralized with 0.1 M glycine in PBS (1 hour) and muscles were blocked with 2% bovine serum albumin (BSA, Sigma), 1% Triton-X (Sigma) in PBS (2 hour). The muscles were subsequently incubated with 0.2 μg/mL mouse anti-SV2 (5ea, developmental studies hybridoma bank) in block ON at 4°C. After 6x 10 minutes washes with PBS, tissue was incubated with 2 μg/mL BTX-488 to visualize AChRs and 2 μg/mL Alexa Fluor 594-conjugated donkey anti-mouse IgG (A21203, Thermo Fisher) in 2% BSA in PBS for 2 hours. After 6x 10 minutes washes in PBS, muscles were mounted in Prolong Gold mounting medium (Thermo Fisher) and stored at 4°C until imaging. Due to technical issues with the staining two ETA muscles (both in the b12 group) and two diaphragm preparations (1x in b12 group and 1x in 13-3B5xb12 group) had to be excluded from further analysis. Consequently, for the ETA muscles the b12 control group was left with only two independent data points. Therefore, statistical analysis was not done on this dataset.

One high resolution Z-stack of a representative part of the muscle was taken using a 20x objective on a SP8 confocal laser-scanning microscope with Las X software (Leica). Z-stacks were converted into maximum projections and further analyzed using ImageJ 1.52n. For the diaphragm images, twenty *en face* NMJs were randomly selected in the 488-BTX-channel and analyzed for intensity and area using a manual threshold. For the analysis of the ETA, thirty enface NMJs were selected, to better capture the variation seen in these preparations. A global threshold was manually determined for the diaphragm and ETA separately. The total AChR signal (intensity X area in the 488-BTX channel) of all analyzed NMJs per image were averaged and used as an n=1 for visualization and further analysis.

NMJ colocalization analysis was conducted on the ETA muscles. All *en face* NMJs were identified in the 488-BTX-channel. An NMJ was excluded if aspecific background in the SV2 channel overlapped with an NMJ. From the remaining NMJs, thirty were randomly selected for colocalization analysis with the EzColocalization plugin for ImageJ (Stauffer et al., 2018). Manually-determined thresholds for each channel were assigned. As a measure of innervation, the fraction of the postsynaptic AChR signal (BTX) that overlaps with the presynaptic signal (SV2), weighted for signal intensity, was quantified for each NMJ with the Mander’s colocalization coefficient M1 (Aaron et al., 2018). In addition, the total presynaptic signal (area x intensity in the SV2 channel) was assessed. All analyzed NMJs per image were averaged and used as an n=1 for visualization.

### Protein A purification of mouse serum

To confirm the integrity of the recombinant antibodies was maintained throughout the *in vivo* experiment, they were purified back from the mouse serum with protein A. To ensure sufficient yield for CE-MS analysis, the mouse sera were pooled per condition. Protein A agarose (Roche) beads were equilibrated with PBS. Pooled serum diluted 1:1 with PBS was incubated with the protein A beads, rotating for 1 hour. Bound protein was eluted with 0.1 M Glycine HCl pH 2.5. Fractions were neutralized with 1:6 1 M Tris-HCl pH 8. Protein content was measured with nanodrop in IgG mode. Protein containing fractions were pooled and immediately processed further for CE-MS as described above.

### Statistics

Data are expressed as mean ± SEM. Comparisons between three or five groups were analyzed using one-way ANOVA with Šidák-corrected comparisons for parametric data. Hanging time (non-parametric data) was analyzed with the Kruskal-Wallis test with Šidák-corrected comparisons. The following comparisons were pre-defined: b12 vs 13-3B5, b12 vs 13-3B5xb12, b12 vs 11-3F6, b12 vs 11-3F6xb12, 13-3B5 vs 13-3B5xb12, 11-3F6 vs 11-3F6xb12, 13-3B5 vs 11-3F6 and 13-3B5xb12 vs 11-3F6xb12. Comparisons between two groups were analyzed using two-tailed unpaired t-test. Measurements from the end-point day was used for statistical analysis by GraphPad Prism (version 8.1.1). Differences were considered significant at p<0.05.

### Study approval

All animal studies were executed with approval of the Dutch national and local animal experiments committees, according the Dutch law and Leiden University guidelines.

## Acknowledgments

We thank Cor Breukel for valuable advice and help on cloning, Steve Burden for the immunoprecipitation antibody and challenging discussions and constructive feedback on our MuSK-related work and Boudewijn Lelieveldt for the advice and help with data visualization. The graphical abstract was created with BioRender.com.

## Funding

The authors are members of the European Reference Network for Rare Neuromuscular Diseases [ERN EURO-NMD]. MH receives financial support from the LUMC (OIO, 2017), Health Holland/TKI Target to B consortium, Prinses Beatrix Spierfonds (W.OR-17.13 and W.OR-19.13) and the Dutch Science Organization NWO (VENI 0915016181 0040). CG receives financial support from the European Commission (Analytics for Biologics; ID 765502).

## Author contributions

DV, JP, CG, PP, MW, EDV, SvdM, JV and MH contributed to the experimental design. DV, JP, CG, YF, RA and RV executed the experiments. DV, JP, CG, RA, RV, EDV, SvdM, JV and MH analyzed and interpreted the data. DV, RV, CG, EDV and MH wrote the manuscript. All authors thoroughly revised and approved the manuscript.

## Notes

**Conflict of interest**: MH, JP, SvdM and JV are co-inventors on two patent applications on MuSK-related research. LUMC, MH, JP, SvdM and JV receive license income from these patents. PP is named inventor on DuoBody-related patents and patent application assigned to Genmab. The authors have no additional financial interest.

### Competing Interest Statement

MH, JP, SvdM and JV are co-inventors on two patent applications on MuSK-related research. LUMC, MH, JP, SvdM and JV receive license income from these patents. PP is named inventor on DuoBody-related patents and patent application assigned to Genmab. The authors have no additional financial interest.

## References

Aalberse, R.C., R. van der Graag, and J. van Leeuwen. 1983. Serologic aspects of IgG4 antibodies. I. Prolonged immunization results in an IgG4-restricted response. J Immunol 130:722–726.

Aaron, J.S., A.B. Taylor, and T.-L. Chew. 2018. Image co-localization – co-occurrence versus correlation. Journal of Cell Science 131:

Adjobimey, T., and A. Hoerauf. 2010. Induction of immunoglobulin G4 in human filariasis: an indicator of immunoregulation. Ann Trop Med Parasitol 104:455–464.

Barbas, C.F., T.A. Collet, W. Amberg, P. Roben, J.M. Binley, D. Hoekstra, D. Cababa, T.M. Jones, R.A. Williamson, G.R. Pilkington, N.L. Haigwood, E. Cabezas, A.C. Satterthwait, I. Sanz, and D.R. Burton. 1993. Molecular Profile of an Antibody Response to HIV-1 as Probed by Combinatorial Libraries. 230:812–823.

Burden, S.J., N. Yumoto, and W. Zhang. 2013. The role of MuSK in synapse formation and neuromuscular disease. Cold Spring Harb Perspect Biol 5:a009167.

Cole, R.N., S.W. Reddel, O.L. Gervásio, and W.D. Phillips. 2008. Anti-MuSK patient antibodies disrupt the mouse neuromuscular junction. Annals of Neurology 63:782–789.

Daveau, M., J. Pavie-Fischer, L. Rivat, C. Rivat, C. Ropartz, H.H. Peter, J.-P. Cesarini, and F.M. Kourlisky. 1977. IgG4 subclass in malignant melanoma. Journal of the National Cancer Institute 58:189–1992.

Eken, T. 1998. Spontaneous Electromyographic Activity in Adult Rat Soleus Muscle. Journal of Neurophysiology 80:365–376.

Evoli, A., P.A. Tonali, L. Padua, M. Lo Monaco, F. Scuderi, A.P. Batocchi, M. Marino, and E. Bartoccioni. 2003. Clinical correlates with anti-MuSK antibodies in generalized seronegative myasthenia gravis. Brain 126:2304–2311.

Fichtner, M.L., C. Vieni, R.L. Redler, L. Kolich, R. Jiang, K. Takata, P. Stathopoulos, P.A. Suarez, R.J. Nowak, S.J. Burden, D.C. Ekiert, and K.C. O’Connor. 2020. Affinity maturation is required for pathogenic monovalent IgG4 autoantibody development in myasthenia gravis. J Exp Med 217:

Fuhrer, C., J.E. Sugiyama, R.G. Taylor, and Z.W. Hall. 1997. Association of muscle-specific kinase MuSK with the acetylcholine receptor in mammalian muscle. The EMBO Journal 16:4951–4960.

Ghazanfari, N., M. Morsch, S.W. Reddel, S.X. Liang, and W.D. Phillips. 2014. Muscle-specific kinase (MuSK) autoantibodies suppress the MuSK pathway and ACh receptor retention at the mouse neuromuscular junction. The Journal of Physiology 592:2881–2897.

Glass, D.J., D.C. Bowen, T.N. Stitt, C. Radziejewski, J. Bruno, T.E. Ryan, D.R. Gies, S. Shah, K. Mattsson, S.J. Burden, P.S. Distefano, D.M. Valenzuela, T.M. Dechiara, and G.D. Yancopoulos. 1996. Agrin acts via a MuSK receptor complex. Cell 85:513–523.

Gstöttner, C., D.L.E. Vergoossen, M. Wuhrer, M.G. Huijbers, and E. Domínguez-Vega. 2020. Sheathless CE-MS as tool for monitoring exchange efficiency and stability of bispecific antibodies. Electrophoresis

Hoch, W., J. McConville, S. Helms, J. Newsom-Davis, A. Melms, and A. Vincent. 2001. Auto-antibodies to the receptor tyrosine kinase MuSK in patients with myasthenia gravis without acetylcholine receptor antibodies. Nature Medicine 7:365–368.

Huijbers, M.G., J.J. Plomp, S.M. van der Maarel, and J.J. Verschuuren. 2018. IgG4-mediated autoimmune diseases: a niche of antibody-mediated disorders. Ann N Y Acad Sci 1413:92–103.

Huijbers, M.G., J.J. Plomp, I.E. van Es, Y.E. Fillié-Grijpma, S. Kamar-Al Majidi, P. Ulrichts, H. de Haard, E. Hofman, S.M. van der Maarel, and J.J. Verschuuren. 2019a. Efgartigimod improves muscle weakness in a mouse model for muscle-specific kinase myasthenia gravis. Experimental Neurology 317:133–143.

Huijbers, M.G., L.A. Querol, E.H. Niks, J.J. Plomp, S.M. van der Maarel, F. Graus, J. Dalmau, I. Illa, and J.J. Verschuuren. 2015. The expanding field of IgG4-mediated neurological autoimmune disorders. European Journal of Neurology 22:1151–1161.

Huijbers, M.G., D.L. Vergoossen, Y.E. Fillie-Grijpma, I.E. van Es, M.T. Koning, L.M. Slot, H. Veelken, J.J. Plomp, S.M. van der Maarel, and J.J. Verschuuren. 2019b. MuSK myasthenia gravis monoclonal antibodies: Valency dictates pathogenicity. Neurol Neuroimmunol Neuroinflamm 6:e547.

Huijbers, M.G., A.F. Vink, E.H. Niks, R.H. Westhuis, E.W. van Zwet, R.H. de Meel, R. Rojas-Garcia, J. Diaz-Manera, J.B. Kuks, R. Klooster, K. Straasheijm, A. Evoli, I. Illa, S.M. van der Maarel, and J.J. Verschuuren. 2016. Longitudinal epitope mapping in MuSK myasthenia gravis: implications for disease severity. J Neuroimmunol 291:82–88.

Huijbers, M.G., W. Zhang, R. Klooster, E.H. Niks, M.B. Friese, K.R. Straasheijm, P.E. Thijssen, H. Vrolijk, J.J. Plomp, P. Vogels, M. Losen, S.M. Van der Maarel, S.J. Burden, and J.J. Verschuuren. 2013. MuSK IgG4 autoantibodies cause myasthenia gravis by inhibiting binding between MuSK and Lrp4. Proc Natl Acad Sci U S A 110:20783–20788.

Ishiguro, T., Y. Sano, S.-I. Komatsu, M. Kamata-Sakurai, A. Kaneko, Y. Kinoshita, H. Shiraiwa, Y. Azuma, T. Tsunenari, Y. Kayukawa, Y. Sonobe, N. Ono, K. Sakata, T. Fujii, Y. Miyazaki, M. Noguchi, M. Endo, A. Harada, W. Frings, E. Fujii, E. Nanba, A. Narita, A. Sakamoto, T. Wakabayashi, H. Konishi, H. Segawa, T. Igawa, T. Tsushima, H. Mutoh, Y. Nishito, M. Takahashi, L. Stewart, E. Elgabry, Y. Kawabe, M. Ishigai, S. Chiba, M. Aoki, K. Hattori, and J. Nezu. 2017. An anti–glypican 3/CD3 bispecific T cell–redirecting antibody for treatment of solid tumors. Science Translational Medicine 9:eaal4291.

Jha, S., K. Xu, T. Maruta, M. Oshima, D.R. Mosier, M.Z. Atassi, and W. Hoch. 2006. Myasthenia gravis induced in mice by immunization with the recombinant extracellular domain of rat muscle-specific kinase (MuSK). J Neuroimmunol 175:107–117.

Karagiannis, P., A.E. Gilbert, D.H. Josephs, N. Ali, T. Dodev, L. Saul, I. Correa, L. Roberts, E. Beddowes, A. Koers, C. Hobbs, S. Ferreira, J.L. Geh, C. Healy, M. Harries, K.M. Acland, P.J. Blower, T. Mitchell, D.J. Fear, J.F. Spicer, K.E. Lacy, F.O. Nestle, and S.N. Karagiannis. 2013. IgG4 subclass antibodies impair antitumor immunity in melanoma. J Clin Invest 123:1457–1474.

Klooster, R., J.J. Plomp, M.G. Huijbers, E.H. Niks, K.R. Straasheijm, F.J. Detmers, P.W. Hermans, K. Sleijpen, A. Verrips, M. Losen, P. Martinez-Martinez, M.H. De Baets, S.M. van der Maarel, and J.J. Verschuuren. 2012. Muscle-specific kinase myasthenia gravis IgG4 autoantibodies cause severe neuromuscular junction dysfunction in mice. Brain 135:1081–1101.

Koneczny, I. 2018. A New Classification System for IgG4 Autoantibodies. Front Immunol 9:97.

Koneczny, I., J. Cossins, P. Waters, D. Beeson, and A. Vincent. 2013. MuSK myasthenia gravis IgG4 disrupts the interaction of LRP4 with MuSK but both IgG4 and IgG1-3 can disperse preformed agrin-independent AChR clusters. PLoS One 8:e80695.

Labrijn, A.F., M.L. Janmaat, J.M. Reichert, and P.W.H.I. Parren. 2019. Bispecific antibodies: a mechanistic review of the pipeline. Nature Reviews Drug Discovery 18:585–608.

Labrijn, A.F., J.I. Meesters, B.E. de Goeij, E.T. van den Bremer, J. Neijssen, M.D. van Kampen, K. Strumane, S. Verploegen, A. Kundu, M.J. Gramer, P.H. van Berkel, J.G. van de Winkel, J. Schuurman, and P.W. Parren. 2013. Efficient generation of stable bispecific IgG1 by controlled Fab-arm exchange. Proc Natl Acad Sci U S A 110:5145–5150.

Labrijn, A.F., J.I. Meesters, P. Priem, R.N. de Jong, E.T. van den Bremer, M.D. van Kampen, A.F. Gerritsen, J. Schuurman, and P.W. Parren. 2014. Controlled Fab-arm exchange for the generation of stable bispecific IgG1. Nat Protoc 9:2450–2463.

Lighaam, L.C., and T. Rispens. 2016. The immunobiology of immunoglobulin G4. Semin Liver Dis 36:200–215.

Liu, H., G. Ponniah, H.-M. Zhang, C. Nowak, A. Neill, N. Gonzalez-Lopez, R. Patel, G. Cheng, A.Z. Kita, and B. Andrien. 2014. In vitro and in vivo modifications of recombinant and human IgG antibodies. mAbs 6:1145–1154.

McConville, J., M.E. Farrugia, D. Beeson, U. Kishore, R. Metcalfe, J. Newsom-Davis, and A. Vincent. 2004. Detection and characterization of MuSK antibodies in seronegative myasthenia gravis. Ann Neurol 55:580–584.

Mori, S., S. Kubo, T. Akiyoshi, S. Yamada, T. Miyazaki, H. Hotta, J. Desaki, M. Kishi, T. Konishi, Y. Nishino, A. Miyazawa, N. Maruyama, and K. Shigemoto. 2012a. Antibodies against muscle-specific kinase impair both presynaptic and postsynaptic functions in a murine model of myasthenia gravis. Am J Pathol 180:798–810.

Mori, S., S. Yamada, S. Kubo, J. Chen, S. Matsuda, M. Shudou, N. Maruyama, and K. Shigemoto. 2012b. Divalent and monovalent autoantibodies cause dysfunction of MuSK by distinct mechanisms in a rabbit model of myasthenia gravis. J Neuroimmunol 244:1–7.

Morsch, M., S.W. Reddel, N. Ghazanfari, K.V. Toyka, and W.D. Phillips. 2012. Muscle specific kinase autoantibodies cause synaptic failure through progressive wastage of postsynaptic acetylcholine receptors. Experimental Neurology 237:286–295.

Nair-Gupta, P., M. Diem, D. Reeves, W. Wang, R. Schulingkamp, K. Sproesser, B. Mattson, B. Heidrich, M. Mendonça, J. Joseph, J. Sendecki, B. Foulk, G. Chu, D. Fink, Q. Jiao, S.-J. Wu, K. Packman, Y. Elsayed, R. Attar, and F. Gaudet. 2020. A novel C2 domain binding CD33xCD3 bispecific antibody with potent T-cell redirection activity against acute myeloid leukemia. Blood Advances 4:906–919.

Neuberger, M.S., and K. Rajewsky. 1981. Activation of mouse complement by monoclonal mouse antibodies. European Journal of Immunology 11:1012–1016.

Niks, E.H., Y. van Leeuwen, M.I. Leite, F.W. Dekker, A.R. Wintzen, P.W. Wirtz, A. Vincent, M.J.D. van Tol, C.M. Jol-van der Zijde, and J.J.G.M. Verschuuren. 2008. Clinical fluctuations in MuSK myasthenia gravis are related to antigen-specific IgG4 instead of IgG1. Journal of Neuroimmunology 195:151–156.

Otsuka, K., M. Ito, B. Ohkawara, A. Masuda, Y. Kawakami, K. Sahashi, H. Nishida, N. Mabuchi, A. Takano, A.G. Engel, and K. Ohno. 2015. Collagen Q and anti-MuSK autoantibody competitively suppress agrin/LRP4/MuSK signaling. Sci Rep 5:13928.

Patel, V., A. Oh, A. Voit, L.G. Sultatos, G.J. Babu, B.A. Wilson, M. Ho, and J.J. McArdle. 2014. Altered active zones, vesicle pools, nerve terminal conductivity, and morphology during experimental MuSK Myasthenia Gravis. PLoS One 9:e110571.

Phillips, W.D., P. Christadoss, M. Losen, A.R. Punga, K. Shigemoto, J. Verschuuren, and A. Vincent. 2015. Guidelines for pre-clinical animal and cellular models of MuSK-myasthenia gravis. Exp Neurol 270:29–40.

Plomp, J.J., M. Morsch, W.D. Phillips, and J.J. Verschuuren. 2015. Electrophysiological analysis of neuromuscular synaptic function in myasthenia gravis patients and animal models. Exp Neurol 270:41–54.

Punga, A.R., S. Lin, F. Oliveri, S. Meinen, and M.A. Rüegg. 2011. Muscle-selective synaptic disassembly and reorganization in MuSK antibody positive MG mice. Experimental Neurology 230:207–217.

Richman, D.P., K. Nishi, S.W. Morell, J.M. Chang, M.J. Ferns, R.L. Wollman, R.A. Maselli, J. Schnier, and M.A. Agius. 2012. Acute severe animal model of anti–muscle-specific kinase Myasthenia. JAMA Neurology 69:453.

Rispens, T., A.M. Davies, P. Ooijevaar-de Heer, S. Absalah, O. Bende, B.J. Sutton, G. Vidarsson, and R.C. Aalberse. 2014. Dynamics of inter-heavy chain interactions in human immunoglobulin G (IgG) subclasses studied by kinetic Fab arm exchange. J Biol Chem 289:6098–6109.

Santos, M.R., C.K. Ratnayake, B. Fonslow, and A. Guttman. 2015. A covalent, cationic polymer coating method for the CESI-MS analysis of intact proteins and polypeptides. In Sciex - Biomarkers Omi.

Schultz, L.D., P.A. Schweitzer, S.W. Christianson, B. Gott, I.B. Schweitzer, B. Tennent, S. McKenna, L. Mobraaten, T.V. Rajan, D.L. Greiner, and E.H. Leiter. 1995. Multiple Defects in Innate and Adaptive Immunologic Function in NOD/LtSz-scid Mice. The Journal of Immunology 154:180–191.

Smith, E.J., K. Olson, L.J. Haber, B. Varghese, P. Duramad, A.D. Tustian, A. Oyejide, J.R. Kirshner, L. Canova, J. Menon, J. Principio, D. Macdonald, J. Kantrowitz, N. Papadopoulos, N. Stahl, G.D. Yancopoulos, G. Thurston, and S. Davis. 2016. A novel, native-format bispecific antibody triggering T-cell killing of B-cells is robustly active in mouse tumor models and cynomolgus monkeys. Scientific Reports 5:17943.

Stauffer, W., H. Sheng, and H.N. Lim. 2018. EzColocalization: An ImageJ plugin for visualizing and measuring colocalization in cells and organisms. Sci Rep 8:15764.

Takata, K., P. Stathopoulos, M. Cao, M. Mané-Damas, M.L. Fichtner, E.S. Benotti, L. Jacobson, P. Waters, S.R. Irani, P. Martinez-Martinez, D. Beeson, M. Losen, A. Vincent, R.J. Nowak, and K.C. O’Connor. 2019. Characterization of pathogenic monoclonal autoantibodies derived from muscle-specific kinase myasthenia gravis patients. JCI Insight 4:

Ulusoy, C., E. Kim, E. Tüzün, R. Huda, V. Yılmaz, K. Poulas, N. Trakas, L. Skriapa, A. Niarchos, R.T. Strait, F.D. Finkelman, S. Turan, P. Zisimopoulou, S. Tzartos, G. Saruhan-Direskeneli, and P. Christadoss. 2014. Preferential production of IgG1, IL-4 and IL-10 in MuSK-immunized mice. Clinical Immunology 151:155–163.

Van Der Neut Kolfschoten, M., J. Schuurman, M. Losen, W.K. Bleeker, P. Martinez-Martinez, E. Vermeulen, T.H. Den Bleker, L. Wiegman, T. Vink, L.A. Aarden, M.H. De Baets, J.G.J. Van De Winkel, R.C. Aalberse, and P.W.H.I. Parren. 2007. Anti-inflammatory activity of human IgG4 antibodies by dynamic Fab arm exchange. Science 317:1554–1557.

Vidarsson, G., G. Dekkers, and T. Rispens. 2014. IgG subclasses and allotypes: from structure to effector functions. Frontiers in Immunology 5:1–17.

Viegas, S., L. Jacobson, P. Waters, J. Cossins, S. Jacob, M.I. Leite, R. Webster, and A. Vincent. 2012. Passive and active immunization models of MuSK-Ab positive myasthenia: electrophysiological evidence for pre and postsynaptic defects. Experimental Neurology 234:506–512.

Xie, M.-H., J. Yuan, C. Adams, and A. Gurney. 1997. Direct demonstration of MuSK involvement in acetylcholine receptor clustering through identification of agonist ScFv. Nature Biotechnology 15:768–771.

Zhou, H., D.J. Glass, G.D. Yancopoulos, and J.R. Sanes. 1999. Distinct domains of Musk mediate its abilities to induce and to associate with postsynaptic specializations. The Journal of Cell Biology 146:1133–1146.

Zhou, L., J. McConville, V. Chaudhry, R.N. Adams, R.L. Skolasky, A. Vincent, and D.B. Drachman. 2004. Clinical comparison of muscle-specific tyrosine kinase (MuSK) antibody-positive and -negative myasthenic patients. Muscle Nerve 30:55–60.

