## Supplementary Material for "Functional monovalency amplifies the pathogenicity of anti-MuSK IgG4 in myasthenia gravis"

**Supplementary Materials**

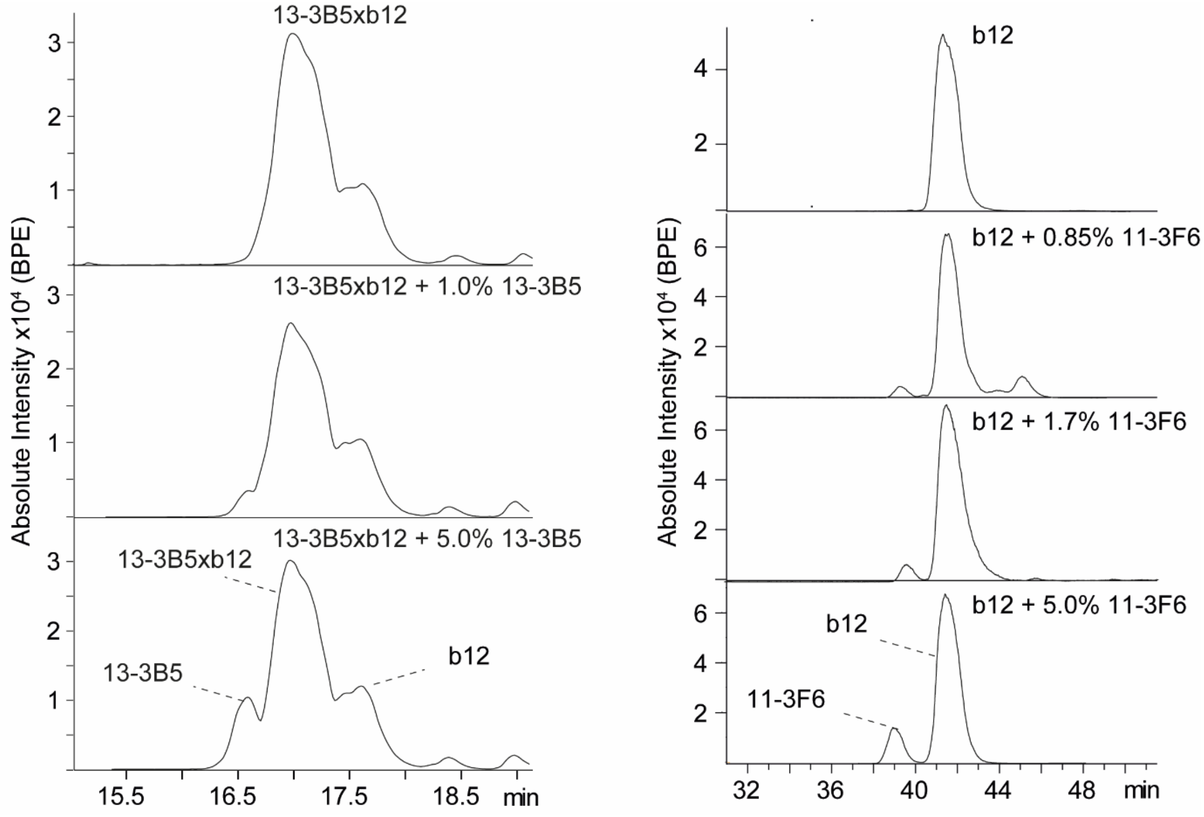


**Supplementary Fig. 1. Detection limit of CE-MS for bivalent 13-3B5 and 11-3F6 is below 0.5%**. To assess the detection limit, antibodies were added in small amounts and analyzed by CE-MS. From the titration curve it is extrapolated that CE-MS permits determination of the bivalent antibodies in relative abundances below 0.5%.

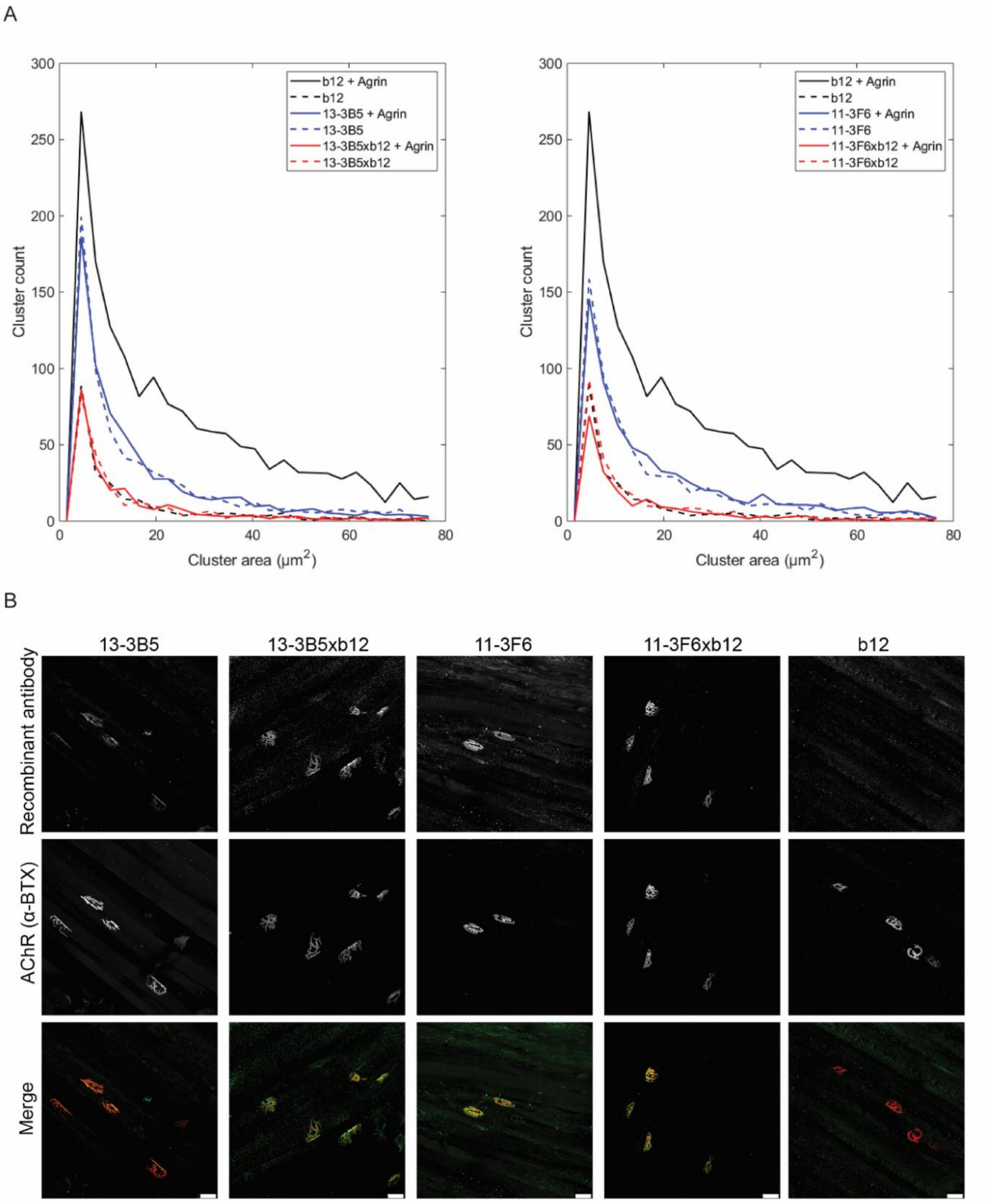


**Supplementary Fig. 2. *In vitro* functional characterization of monovalent and bivalent IgG4 MuSK antibodies**. (A) Distribution of AChR clusters based on cluster size (cutoff 3 μm$^2$) revealed that bivalent 13-3B5 induced more small clusters compared to bivalent 11-3F6 in C2C12 differentiated myotubes. The effect of the bivalent and monovalent anti-MuSK clones was independent of agrin. Distributions represent mean of three independent experiments. (B) Bivalent and monovalent MuSK antibodies bind NMJs of NOD/SCID mice. The control b12 antibody did not show binding. α-BTX = alpha-bungarotoxin. Scale bar 25 μm.

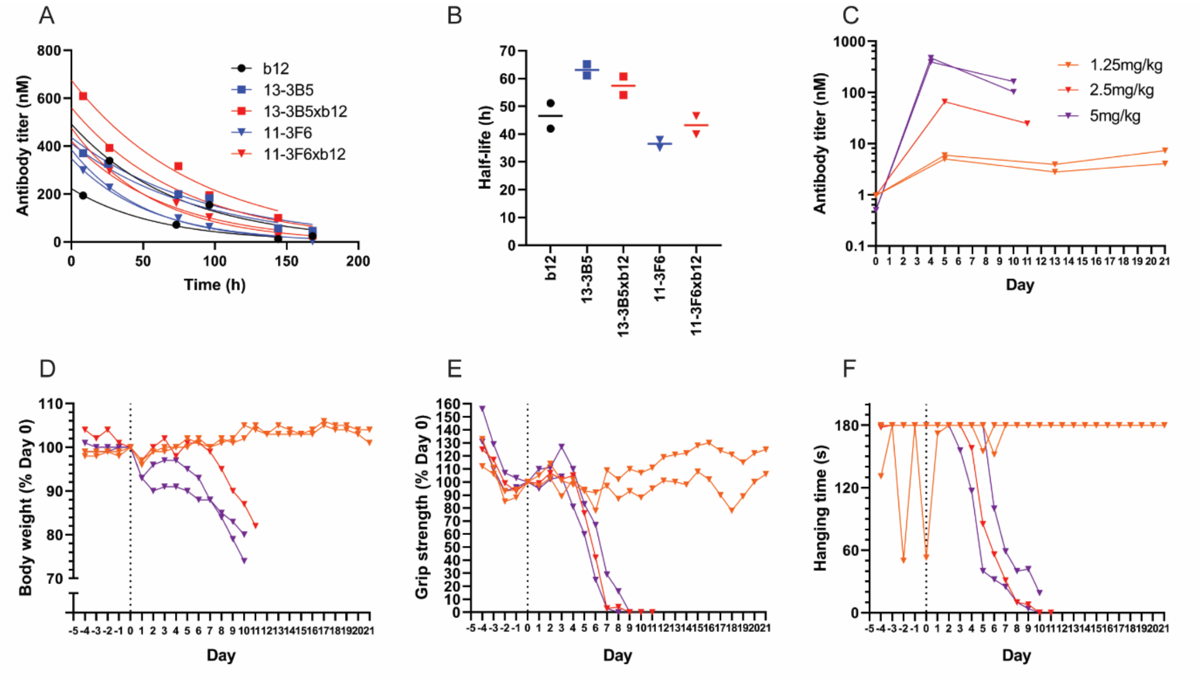


**Supplementary Fig. 3. Half-life and *in vivo* dose-finding**. (A and B) Serum antibody titer was assessed at different times after a single i.p. dose of 5 mg/kg recombinant antibody with antigen-specific ELISA (n=2). The half-life was calculated by fitting a one-phase exponential decay function with the plateau constrained at zero. Half-life ranged between 38-63 hours, depending on the antibody. (C) MuSK antibody titers of NOD/SCID mice injected with different doses of 11-3F6xb12 every 3-4 days. (D) Body weight, (E) grip strength and (F) inverted mesh hanging time revealed that 2.5 mg/kg every 3-4 days was the minimal dose required to cause progressive phenotypical myasthenia (n=1-2). Individual data are presented.

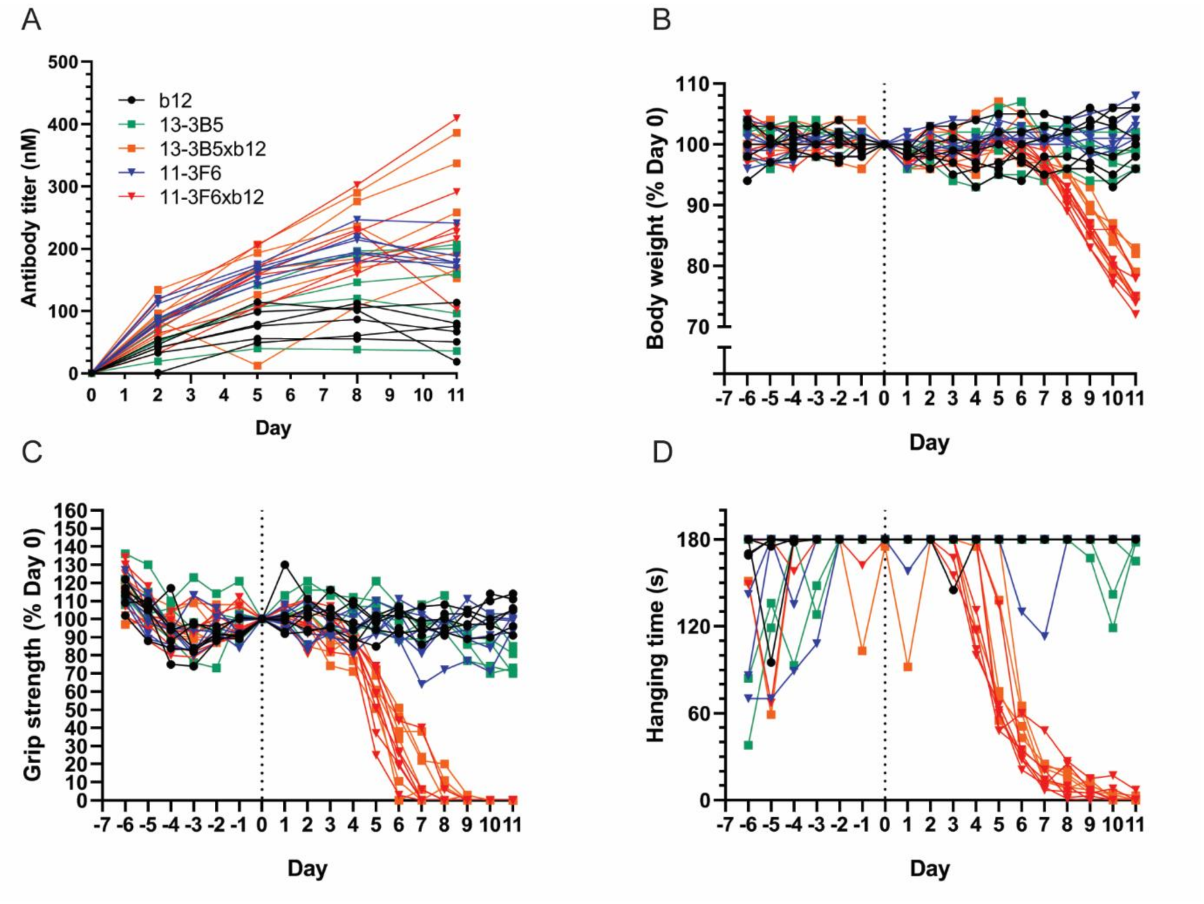


**Supplementary Fig. 4. Individual mouse data of *in vivo* parameters**. Data trajectories of Figure 3 visualized per mouse for (A) antibody titer, (B) body weight, (C) grip strength and (D) inverted mesh hanging time. 11-3F6 and 11-3F6xb12 n=6, 13-3B5 n=5, 13-3B5xb12 and b12 n=6 (hanging time n=5).

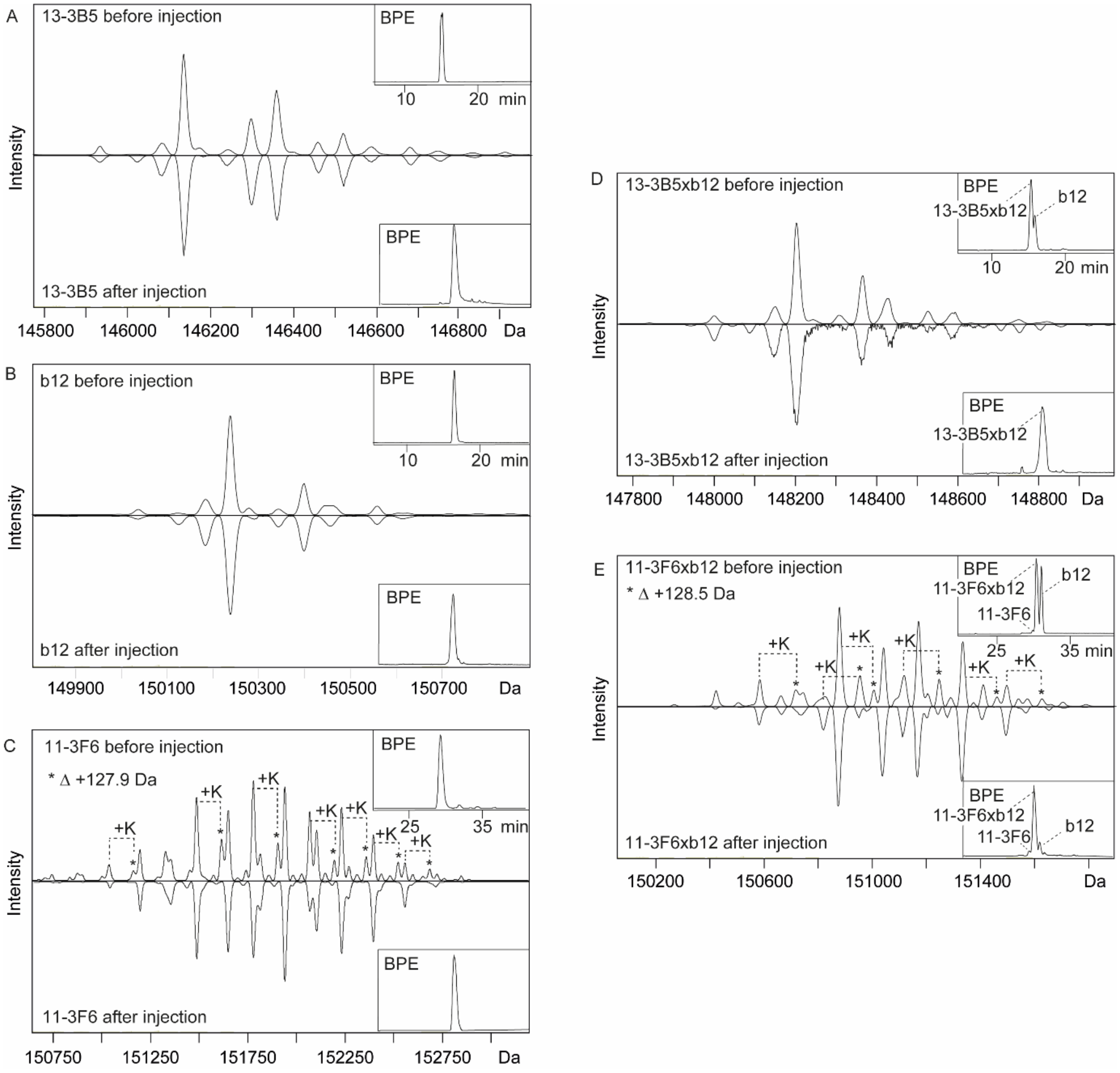


**Supplementary Fig. 5. Recombinant bivalent and monovalent antibodies are stable *in vivo*.** Deconvoluted mass spectra and base electropherograms (BPE) of bivalent and monovalent antibodies before (upper trace) and after (lower trace) injection in mice showed stability of the antibodies *in vivo*. (A) Bivalent 13-3B5, (B) bivalent b12, (C) bivalent 11-3F6, (D) monovalent 13-3B5xb12 and (E) monovalent 11-3F6xb12. 11-3F6 (C, upper trace) and 11-3F6xb12 (E, upper trace) contained variants with an additional lysine (K) at the C-terminus of the Fc tail, indicated by (*). C-terminal lysines on IgG Fc are known to be clipped in the circulation (Liu et al., 2014) and could consequently not be observed in the mass spectra of 11-3F6 (C, lower trace) and 11-3F6xb12 (E, lower trace) purified from mouse serum.

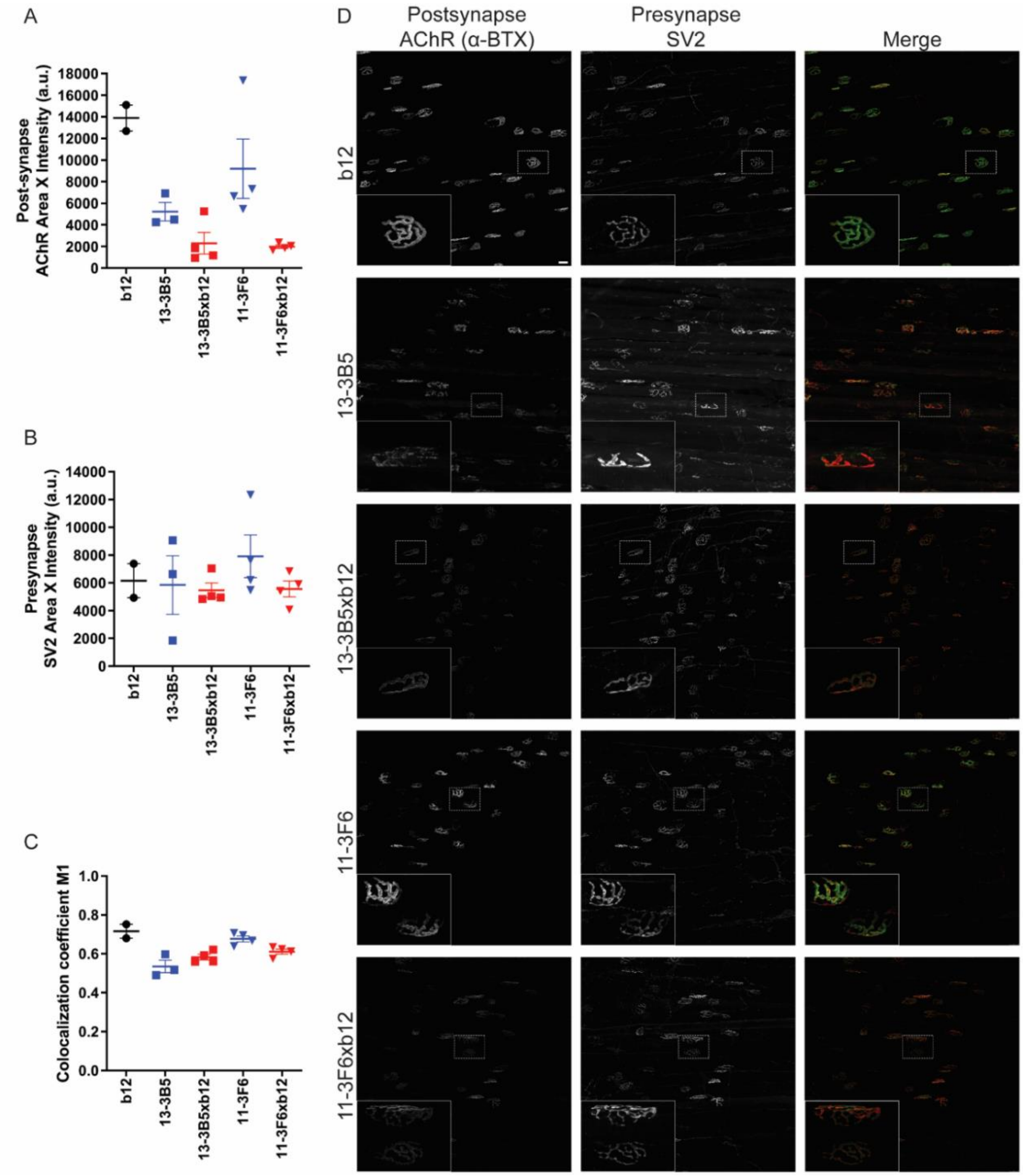


**Supplementary Fig. 6. Altered NMJ morphology caused by bivalent and monovalent MuSK antibodies seems to be limited to the postsynapse**. AChR staining (AF488-BTX) marks the postsynaptic NMJ area and synaptic vesicle protein 2 (SV2) staining marks the presynaptic NMJ area. Thirty randomly selected NMJs per epitrochleoanconeus (ETA) muscle were analyzed and averaged. (A) Exposure to bivalent or monovalent MuSK antibodies resulted in, on average, less postsynaptic signal. (B) Presynaptic morphology does not seem to be affected by exposure to monovalent or bivalent MuSK antibodies. (C) NMJ innervation by the motor neuron was assessed by the colocalization coefficient M1, which is an intensity weighted co-occurrence coefficient of presynaptic signal (SV2) with postsynaptic signal (AChR) compared to the total postsynaptic area. Monovalent MuSK antibodies and bivalent 13-3B5 seem to slightly reduce the overlap between the pre- and postsynapse. (D) Representative maximum projections with insets per condition. In the merged picture green = AChR, red = SV2. Scalebar = 25 µm. 11-3F6, 11-3F6xb12 and 13-3B5xb12 n=4, 13-3B5 n=3 and b12 n=2. Data represents mean ± SEM.

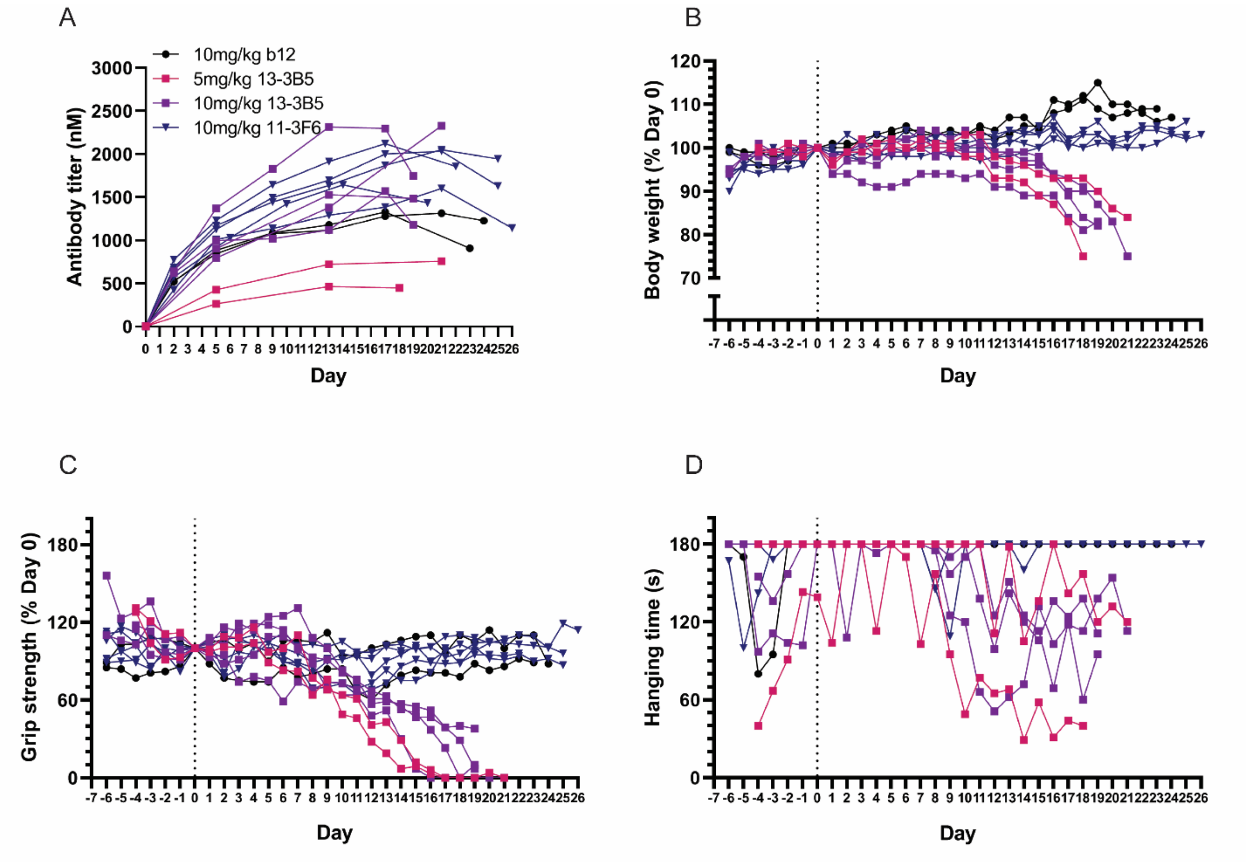


**Supplementary Fig. 7. Individual mouse data of *in vivo* parameters**. Data trajectories of Figure 5 visualized per mouse for (A) antibody titer, (B) body weight, (C) grip strength and (D) inverted mesh hanging time. 10mg/kg b12 n=2, 5mg/kg 13-3B5 n=2, 10mg/kg 13-3B5 n=4 (hanging time n=3), 10mg/kg 11-3F6 n=5.

**Supplementary Table 1: Predicted molecular characteristics of recombinant antibodies, based on amino acid sequence (glycosylation was not considered).**

| Recombinant antibody | Mass (kDa) | Molar extinction coefficient ($x10^3$ $M^{-1}cm^{-1}$) | Absorption coefficient |
|---|---|---|---|
| **b12** | 147.67 | 218.42 | 1.48 |
| **13-3B5** | 143.57 | 215.44 | 1.50 |
| **13-3B5xb12** | 145.62 | 216.93 | 1.49 |
| **11-3F6** | 144.76 | 196.42 | 1.36 |
| **11-3F6xb12** | 146.22 | 207.42 | 1.42 |
